# Atlas of nascent RNA transcripts reveals enhancer to gene linkages

**DOI:** 10.1101/2023.12.07.570626

**Authors:** Rutendo F. Sigauke, Lynn Sanford, Zachary L. Maas, Taylor Jones, Jacob T. Stanley, Hope A. Townsend, Mary A. Allen, Robin D. Dowell

**Author notes:** Contributing authors.

## Abstract

Gene transcription is controlled and modulated by regulatory regions, including enhancers and promoters. These regions are abundant in unstable, non-coding bidirectional transcription. Using nascent RNA transcription data across hundreds of human samples, we identified over 800,000 regions containing bidirectional transcription. We then identify highly correlated transcription between bidirectional and gene regions. The identified correlated pairs, a bidirectional region and a gene, are enriched for disease associated SNPs and often supported by independent 3D data. We present these resources as an SQL database which serves as a resource for future studies into gene regulation, enhancer associated RNAs, and transcription factors.

## Introduction

Transcription is a regulated process that is critical for cellular identity, differentiation, and response to the environment. Nascent transcription assays provide insight into transcription by measuring RNAs prior to their maturation into messenger RNA. In particular, run-on assays such as global run-on sequencing (GRO-seq) and precision run-on sequencing (PRO-seq) measure RNA as it is being produced by incorporating a marked nucleotide and selectively precipitating the labeled RNA. In this manner, the activity of cellular polymerases can be precisely measured.

Mammalian transcription initiation is predominantly bidirectional, with two oppositely oriented distinct transcription start sites in close proximity[1, 2]. These bidirectional signatures are observed at not only protein-coding genes but also transcribed regulatory regions (TREs)[2–4]. At genes, the upstream antisense RNA (uaRNA) is also referred to as a promoter upstream transcript (PROMPTs)[5–7]. At regulatory regions, the two transcripts are often referred to as enhancer associated RNAs (eRNAs)[7–10]. In every case, these non-gene transcripts are lowly transcribed, unstable, and not annotated. Hence numerous methods have been developed to identify sites of bidirectional transcription directly from run-on data[4, 11–14].

The function of these transcripts remains hotly debated, but regardless of their function, they have been shown to serve as excellent markers of regulation within the local genomic context. Not all instances of transcription factor binding result in altered gene regulation, however signatures of RNA polymerase activity near transcription factor binding sites effectively reflects the active subset of binding events[15–18]. In support of this notion, changes in bidirectional transcription activity (locations and levels) can be utilized to infer changes in transcription factor activity between two conditions[8, 18–24].

Such activity at regulatory regions can be essential for understanding changes in transcription at associated target genes. However, while some TREs have been definitively linked to target genes, the majority of these linkages are unknown. Enhancer regions are more distal to genes than promoters, which complicates the process of correctly assigning them to their targets. Yet transcription levels at regulatory regions clearly correlate with transcription levels at their associated target gene[18, 25]. In fact, correlated transcription has recently been used in small collections of data to link enhancers to their target genes[26, 27]. This suggests that patterns of correlation maintained over a large collection of nascent transcription data would identify enhancer to target gene pairings, which could then be used to decipher regulatory variation and refine our understanding of gene transcription regulation.

We sought to collect a repository of published run-on sequencing data from which we could catalog and characterize sites of bidirectional transcription. In total we collected thousands of samples from the sequencing archives from which we annotated hundreds of thousands of sites of bidirectional transcription. The majority of these sites did not reside at promoters and were either cell type or tissue specific. Finally, we link sites of bidirectional transcription to target genes, showing that many highly correlated pairs are supported by known enhancer-gene linkages. Our repository will serve as valuable resource for future studies into transcriptional regulation.

## Results

### A repository of run-on nascent RNA data

We began by assembling a large repository of previously published nascent transcription data sets (Figure 1). To this end, nascent RNA sequencing experiments were manually curated from the Gene Expression Omnibus (GEO) [28, 29] and the NIH Sequence Read Archive (SRA) [30]. We excluded metabolic labeling techniques as recovery of transcribed regulatory elements is highly sensitive to the length of incubation with the marked nucleotide. Metadata details such as organism, cell type, protocol used, library preparation, treatment type/conditions, and replicate information were collected for all samples from their associated database information and/or publication (See Supplementary Table 1). This metadata was collected into a MySQL database (hereafter DBNascent) where all treatment condition times were annotated in reference to the time of cell harvest. Raw fastq files were processed through standardized Nextflow pipelines (Figure 1A) that include mapping, quality control, and identifying regions of bidirectional transcription. In all cases, technical replicate fastq files were combined for downstream analysis.

**Fig. 1.**
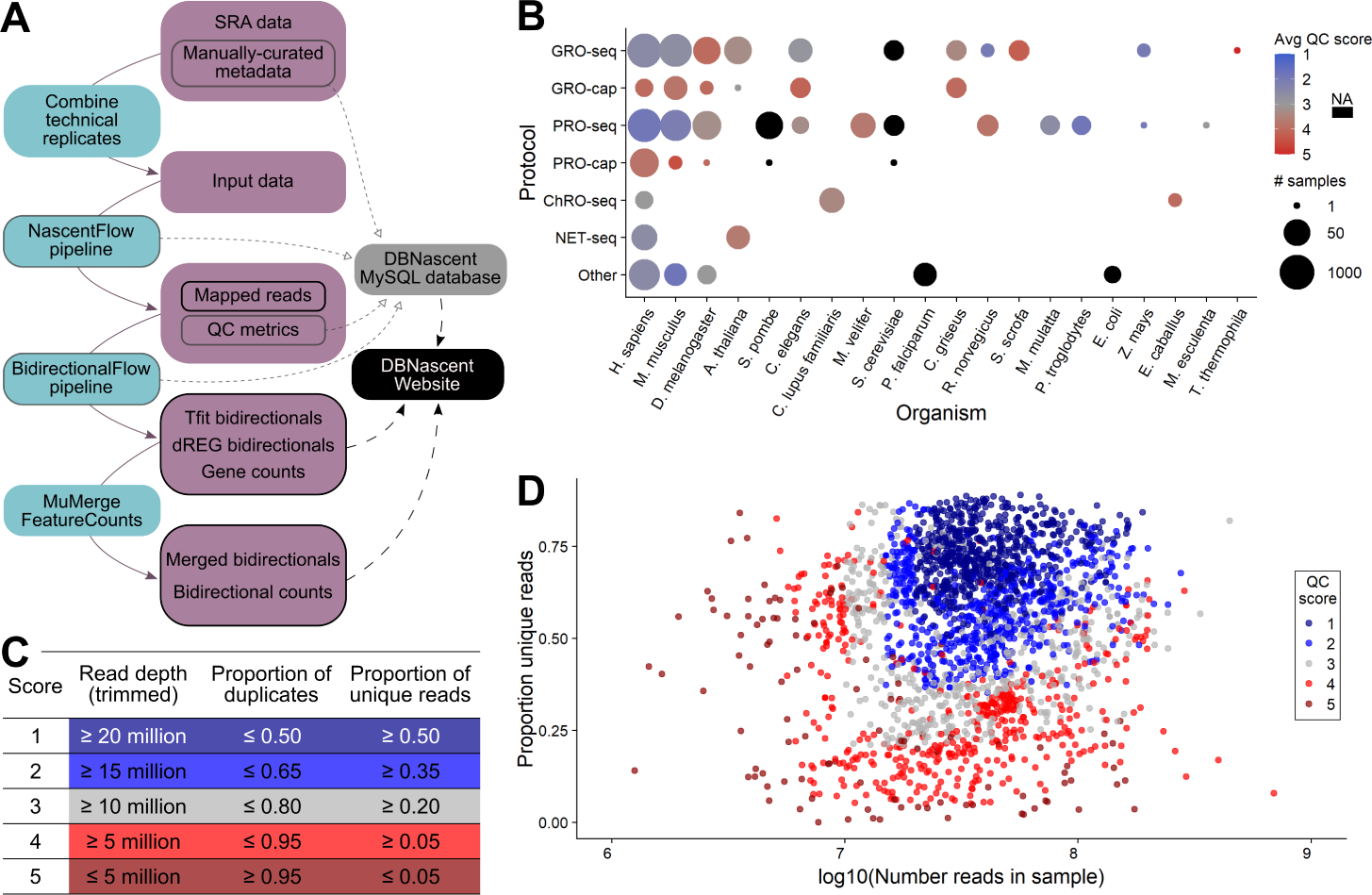
Overview of DBNascent. (A) Data were derived from Sequence Read Archive fastq files and manually curated metadata. Technical replicate fastq files were combined, then data were processed to obtain metrics on quality, bidirectional regions, and read counts. Metadata, quality control metrics, and software version information from the pipeline were accumulated into a MySQL database. The DBNascent website (nascent.colorado.edu) draws from the MySQL database as well as processed analysis files for visualization and region-specific read counts. (B) Samples in DBNascent were derived from twenty different organisms and multiple different protocols. All species with genomes less than 25 Mb were not described well by the calculated QC score and thus are represented as black (NA). (C) Thresholds for calculation of the QC score, tuned for mammalian samples. (D) Complexity (yaxis) versus read depth (x-axis) of human and mouse samples. Two very low read depth samples have been omitted for the sake of visualization.

In total, 3,638 raw samples from the NIH Sequence Read Archive (SRA) were combined into 2,880 biological samples across 20 organisms, collected from 287 projects, which consisted of either journal articles or Gene Expression Omnibus (GEO) datasets (Figure 1B). The samples were subjected to extensive quality control (QC), from which we developed a QC ranking metric based on both read depth and complexity (Figure 1C-D). We used this metric extensively as a filtering mechanism, and most downstream analyses using high quality samples with a QC score of 1-3, unless specified otherwise. As run-on assays necessarily depend on a pull down step involving antibodies, we also sought to assess the extent of nascent RNA enrichment. To this end, we developed an additional score to identify samples that exhibited patterns of nuclear run-on (NRO) sequencing, which could then be used as another potential filtering metric (Supplementary Figure 1).

Of the 2,880 samples in DBNascent, the vast majority (2,387) were derived from either human or mouse cells (Figure 1B), and these were exclusively used for downstream analysis, e.g. identifying bidirectional regions. Samples were distributed across 19 and 10 tissues from human and mouse, respectively. In both organisms, samples were mainly collected from cell lines or cultured primary cells (Supplementary Figure 2). Additionally, a principle component analysis on high quality human samples indicates that samples cluster predominantly by tissue of origin rather than quality score, indicating that differences in the data reflect underlying biological signal more than technical variation (Supplementary Figure 3).

### Bidirectional regions in DBNascent overlap cis-regulatory elements

Nuclear run-on assays, such as GRO-seq and PRO-seq, give readout of transcription from all cellular RNA polymerases. Consequently, they recover signal at both coding and non-coding regions, much of which is not annotated. Two methods for identifying transcribed regulatory regions are Tfit and dREG[4, 11, 12]. Tfit uses a mathematical model of RNA polymerase II to identify sites of polymerase loading and initiation, the majority of which are bidirectional. In contrast, dREG uses an unsupervised support vector machine approach to identify transcribed regulatory elements (TREs), most of which show bidirectional transcription. The two approaches are thus quite distinct and complementary, but both seek to identify sites of bidirectional transcription directly from the data.

As the two methods have distinct strengths and weaknesses, we combined the results of both methods to identify sites of bidirectional transcription (see Methods for complete details). For each of the 1,638 human and 750 mouse samples analyzed, there were on average ~ 25,000 bidirectional regions identified by Tfit and ~ 18,500 by dREG (Supplementary Figure 4A). Bidirectional calls were then combined using a modified version of *muMerge* (version 1.1.0) (see Methods Section) [19]. The merging strategy was performed in a hierarchical manner, merging across experiments first, then across cell type and finally between the bidirectional calling methods (Figure 2A). Since the resolution of Tfit calls at RNA polymerase initiation (typically the center region of bidirectional transcription) is better than dREG [14], the coordinates of Tfit calls were used when the two callers overlap (Supplementary Figure 4B-C). Called regions were filtered to retain high quality regions, based on the data’s QC score (Supplementary Figure 5).

**Fig. 2.**
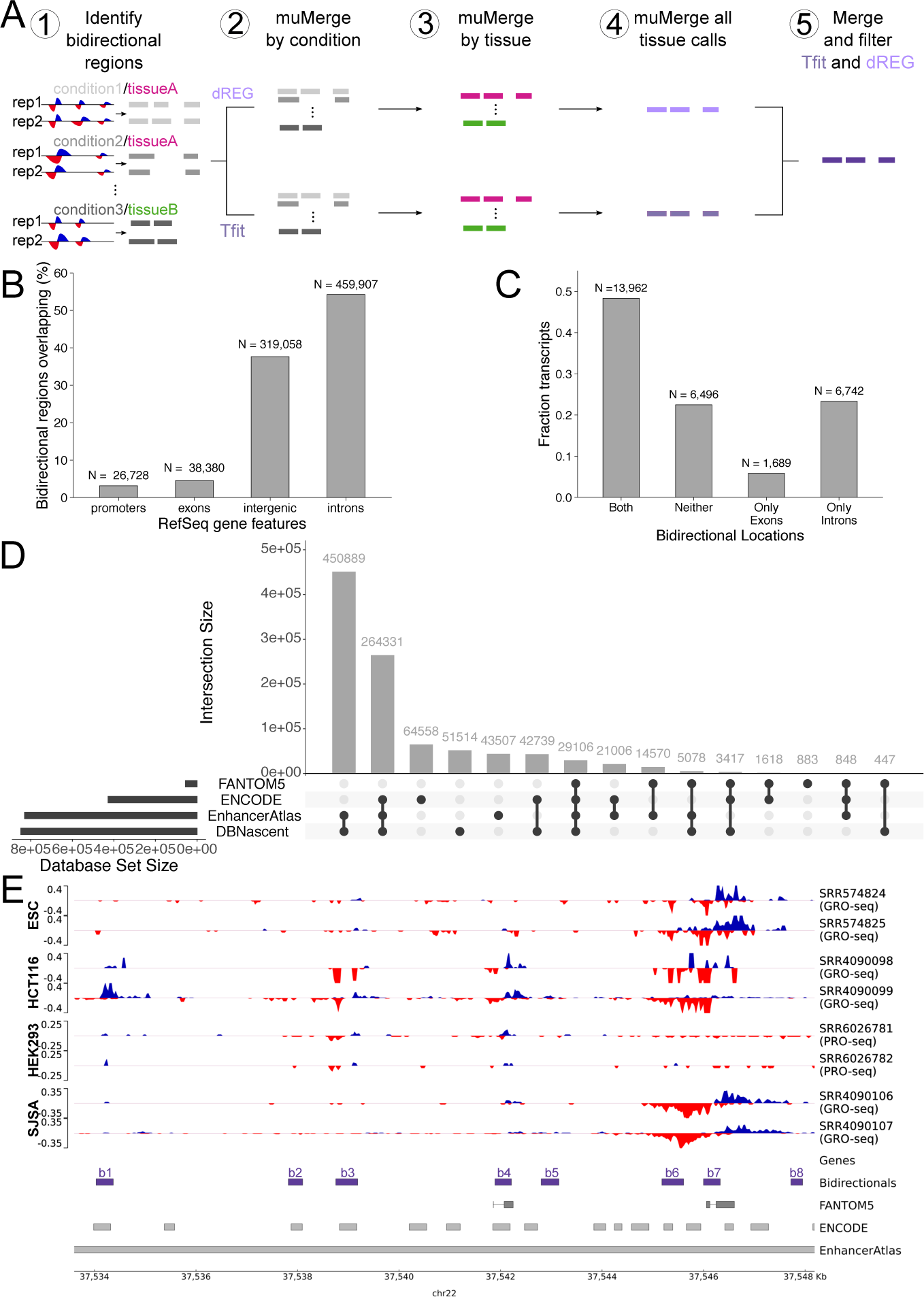
Identification and characterization of bidirectional regions. (A) Schematic showing how bidirectional regions were identified and merged to give a consensus annotation set. Briefly, (1) First, bidirectional regions were inferred from nuclear run-on coverage data in each sample. (2) In a given experiment, regions were combined using muMerge[19] based on treatment conditions for both Tfit[11] and dREG[4, 12] called bidirectional regions. (3) Tfit and dREG calls were combined by muMerge based on cell/tissue type. (4) Master call lists were then obtained by muMerge and (5) combined and filtered. (B) Overlap between bidirectional regions and RefSeq hg38 gene features (exons, introns, promoters and intergenic regions). (C) Fraction of RefSeq annotated genes with bidirectional regions overlapping their introns and/or exons. (D) Overlap between bidirectional regions (DBNascent) and cis-regulatory elements from other databases (ENCODE[33–36], FANTOM5[3, 25, 31, 32] and EnhancerAtlas[37]). (E) An example region (chr22:37,533,600-37,548,188) showing mapped read coverage for tw_1_o_1_replicates each of four cell lines (ESCs, HCT116, HEK293, and SJSAs) along with bidirectional region calls (purple) and regulatory regions identified by FANTOM5, ENCODE and EnhancerAtlas. Importantly, each inferred bidirectional region has a distinct cell type and tissue specific transcription profile. b1: HCT116 only, b3, b4: HCT116 and SJSA, b7: ESC, HCT116 and SJSA, b2, b5, b6 and b8: not in these four cell lines; blue: positive strand, red: negative strand.

Genome wide, 847,521 unique bidirectional calls were obtained across all human data sets and 680,735 in mouse. Bidirectional regions, as expected, are generally much shorter than genes (Supplementary Figure 6) and are similarly distributed across the genome (Supplementary Figures 7 and 8). The majority of bidirectional regions overlap non-coding regions, while a smaller percentage are in exons (Figure 2B and Supplementary Figure 9A). Characterizing the number of human genes with bidirectional regions, we find that about 80% have a bidirectional call in their promoter region, and the transcripts without a bidirectional call at their TSS were not transcribed (Supplementary Tables 2 and 3). Outside the promoter region, bidirectional regions are found uniformly across annotated transcripts in both mouse and human (Supplementary Figure 10. Bidirectional regions within the gene are mostly intronic, with many overlapping the boundaries with an exon (Figure 2C and Supplementary Figure 9B).

To assess the quality of our called regions, we next compared our bidirectional calls to annotated cis-regulatory elements from ENCODE, EnhancerAtlas and FANTOM5, as these resources annotate regulatory regions using a variety of techniques [3, 25, 31–37] (Figure 2D and Supplementary Figure 11). ENCODE offers a large characterization of cis-regulatory elements based on histone, DNase and CTCF signal [33, 34]. While the FANTOM5 project identifies sites of transcription initiation primarily using CAGE (Cap Analysis of Gene Expression) data [32]. Lastly, EnhancerAtlas aims to combine assorted genomic data, including ENCODE and FANTOM5 as well as nascent RNA sequencing data [25, 31, 37]. Overall, about 40% to 60% of the cis-regulatory elements in these data resources are found in DBNascent (Supplementary Figures 11A and B). Interestingly, 29,106 human and 21,999 mouse bidirectional regions are contained in all three databases (Figure 2D and Supplementary Figure 11C). In general, we found a greater overlap between bidirectional regions and EnhancerAtlas regions. However, upon closer examination we noticed that EnhancerAtlas regions tend to be wider compared to all the other database regions therefore yielding greater overlaps (Figure 2E and Supplementary Figure 12). Notably, EnhancerAtlas includes nascent RNA data and RNA polmerase II ChIP seq in its construction, which may contribute to both the observed region length and the overlap with our called bidirectional regions. In conclusion, we recover a large fraction of the previously annotated cis-regulatory elements, despite having data from far fewer tissues than was used in these databases. Regulatory regions have also been identified based on large scale genome-wide association studies. In particular, the GTEx consortium examined genome variation for its ability to influence expression levels [38]. As sites of bidirectional transcription are often genetic enhancers, we next considered to what extent our bidirectional calls overlap with GTEx identified variation. While only a small number of GTEx variation resides within our bidirectional regions (Supplemental Figure 11A), we found that bidirectional regions showed a higher odds for containing significant expression quantitative trait loci (eQTL) variants compared to non-significant variants (Supplementary Figure 13) [38]. This further supports previous work showing an enrichment of eQTLs in enhancer regions [4].

### Tissue specificity of transcription

We next sought to determine how transcription levels varied across different types of transcribed regions. Within a representative high-quality dataset [39], promoter bidirectional regions were most likely to be highly transcribed, followed by both coding and annotated noncoding genes (Figure 3A). Collectively, the exonic, intronic, and intergenic bidirectional regions (non-promoter bidirectional regions), which tend to be enhancers, are much more lowly expressed. This pattern held true across all 741 human samples, where coding genes and promoter bidirectional regions were more highly transcribed with less variability across samples than non-promoter bidirectional regions or noncoding genes (Figure 3B).

**Fig. 3.**
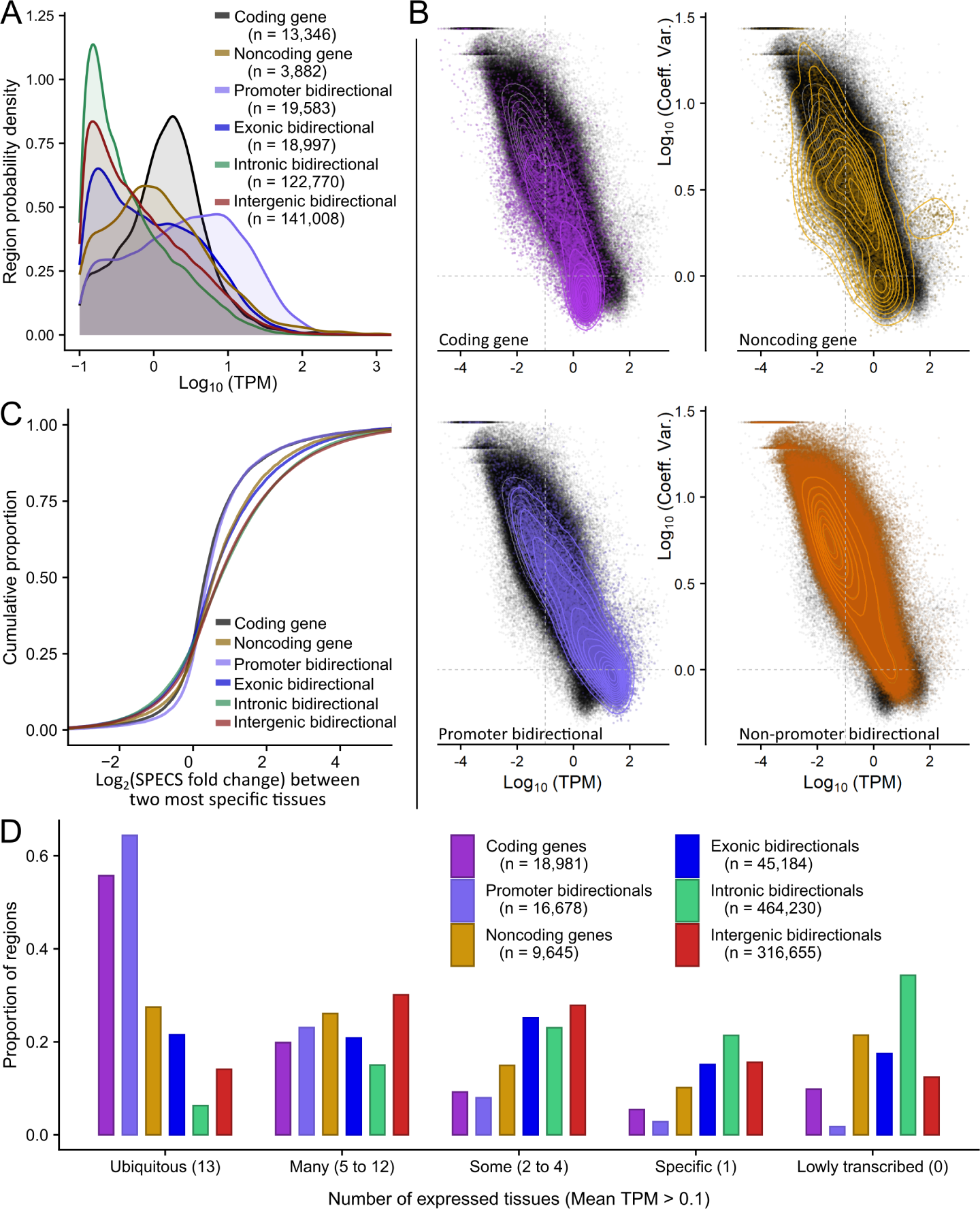
Variation in transcription levels and tissue specificity across annotation types. (A) Distribution of average TPMs (x-axis) for different classes of regions across replicate high-quality MCF7 datasets (SRR5227979 and SRR5227980). Number of regions (*n*) with average TPM *>* 0.1. (B) Across high quality human samples, the coefficient of variation (y-axis) of each region class compared to transcription (x-axis, log(TPM)). Black points and gray density contours display all regions in all plots, overlaid by region-specific colored points and density contours. ’Non-promoter bidirectionals’ includes intronic, exonic, and intergenic regions. (C) Cumulative distribution of fold changes between the tissue with the highest SPECS score and the tissue with the second highest SPECS score for each region class. This strategy is adapted from Everaert et al. 2020[40]. (D) Number of tissues (x-axis) in which a region is transcribed, by class of region. ’Lowly transcribed’ refers to regions that failed to reach the TPM threshold of 0.1 in any tissue.

Due to the consistent trends of the magnitude of bidirectional transcription within given region classes, we investigated the tissue specificity for these classes. We limited this investigation to samples with a QC score of 1-3 that were derived from unique tissues at least 5 samples in the database. As the number of samples in each tissue varied widely, we chose to assess tissue specificity with the SPECS score [40], which can accommodate uneven sample size across groups. The SPECS score ranges from 0 (indicating depletion) to 1 (indicating enrichment), with a ubiquitously transcribed gene scoring around 0.5. Considering all genes and bidirectional regions, the distribution of SPECS scores showed a larger proportion of bidirectional regions having lower SPECS scores, indicating they are more likely to be show higher expression in a limited set of tissues and low (or no) expression across all others (Supplementary Figure 14). For a given tissue, both genes and bidirectional regions had similar trends of high SPEC scores, with umbilical cord, prostate, and uterine samples containing the highest numbers of tissue specific genes and bidirectional regions (Supplementary Figure 15).

We next assessed the change in transcription between the most specific tissue (highest SPECS score) and next highest scoring tissue (Figure 3C). The resulting fold change should be large for each transcript which is transcribed primarily in a single tissue. Interestingly, we observed a skew towards higher fold changes for non-promoter bidirectional regions as compared to genes. Promoter bidirectional calls showed a pattern indistinguishable from coding genes, whereas noncoding genes seemed to show more tissue specificity than coding genes, in line with previous work [41]. Within nonpromoter bidirectional regions, those overlapping with exons were less tissue specific than intergenic or intronic bidirectional calls, likely due to some exonic bidirectional regions toward the 5’ end of genes having spillover transcription signal from promoter regions.

The SPECS score analysis suggests that non-promoter bidirectional calls, primarily associated with enhancers, are the most tissue specific transcripts. To further evaluate this claim, for each region type we quantified 1) the number of tissues in which it was transcribed and 2) the variation of that transcription level. (Supplementary Figure 16, Supplementary Video 1). In all region classes, ubiquitously transcribed regions (transcribed in all 13 tissues) showed much less variation in transcription levels than tissue-specific regions (only present in one tissue). We further investigated the proportion of regions transcribed across the tissues analyzed (Figure 3D). By this measure, coding regions and promoter bidirectional regions are most likely to be ubiquitously transcribed, whereas intronic and intergenic bidirectional regions are most likely to show tissue specific transcription, consistent with previous reports[26]. Thus intronic and intergenic bidirectional regions associated with enhancers are transcribed and active in a small range of tissues compared to coding and noncoding genes.

### Correlation analysis to identify putative bidirectional and gene pairs

Various methods have been developed to link enhancers to target genes with genomic sequencing data [25, 42–44]. Initially, the closest gene approach was primarily used for assigning enhancers (or ChIP sites) to target genes. However, the closest gene is not always accurate, as indicated by 3D information [45]. Prior work on nascent transcription showed that enhancers and their known target genes – as determined by 3D data – have correlated transcription levels[7, 26, 27, 46]. Therefore we sought to determine whether correlation across the collection was sufficient to identify enhancer to target gene linkages.

To this end, we calculated pairwise gene and bidirectional correlations within each chromosome and identified highly correlated pairs in a tissue specific manner for human samples (Supplementary Figure 17) (See Methods). In a collection of bidirectional regions and genes we found 1,094,246 unique pairs where the absolute Pearson correlation coefficient (PCC) was greater than 0.6, and the adjusted p-value was less than 0.01. Most pairs are on chromosome 1 (Supplementary Figure 18A), consistent with its large size (~248 Mb) and high gene density.

While not a constraint of the approach, we found that most bidirectional regions within the pair were close to the gene TSS (Supplementary Figure 18B). Across these pairs, the median number of bidirectional regions assigned to a gene was 42, indicating that many bidirectional regions may factor into tuning the transcription level of a given gene. However, within the context of a single tissue we observe fewer bidirectional regions linked to a gene (median = 2-8; Supplemental Figure 19). This estimate is on par with other estimates of number of enhancers linked to a gene[47–54]. In the other direction, the median number of genes assigned to a bidirectional transcript across all tissues was four, implying that a single bidirectional has only a few potential gene targets (Supplementary Figures 18C and D). As before, this decreases in a tissue specific context (median = 1-3; Supplementary Figure 20).

When assessing the number of tissues that support a pair, approximately 35% of pairs were supported by two or more tissues (Figure 4A and Supplementary Figure 21A). In total, 82.7% of genes are linked to a bidirectional transcript, while only 21.15% of the bidirectional regions have links (Supplementary Figures 22A and B). Interestingly, there was no relationship between SPECS scores and the number of tissues supporting a gene and bidirectional link (Supplementary Figures 23A and B). We next sought to determine whether our correlated pairs were enriched for biologically meaningful pairs. To this end, we took a randomization strategy. We reasoned most biologically meaningful correlation would break down if the data were selected randomly from all bidirectional regions not on the current chromosome. Thus we shuffled the data associated with each bidirectional position, sampling the vector of transcription profiles from all other chromosomes. We then compared assigned pairs from the within chromosome comparisons to cross chromosome comparisons, finding that within chromosome comparisons had far more assigned pairs (Figure 4B), suggesting our assigned pairs contain real signal beyond random correlations. Notably, this randomization is imperfect, as we would retain some real correlation signal when the randomly selected bidirectional and current bidirectional shared an upstream regulator. Thus our randomization likely underestimates the proportion of real biological induced correlation relative to spurious correlations.

**Fig. 4.**
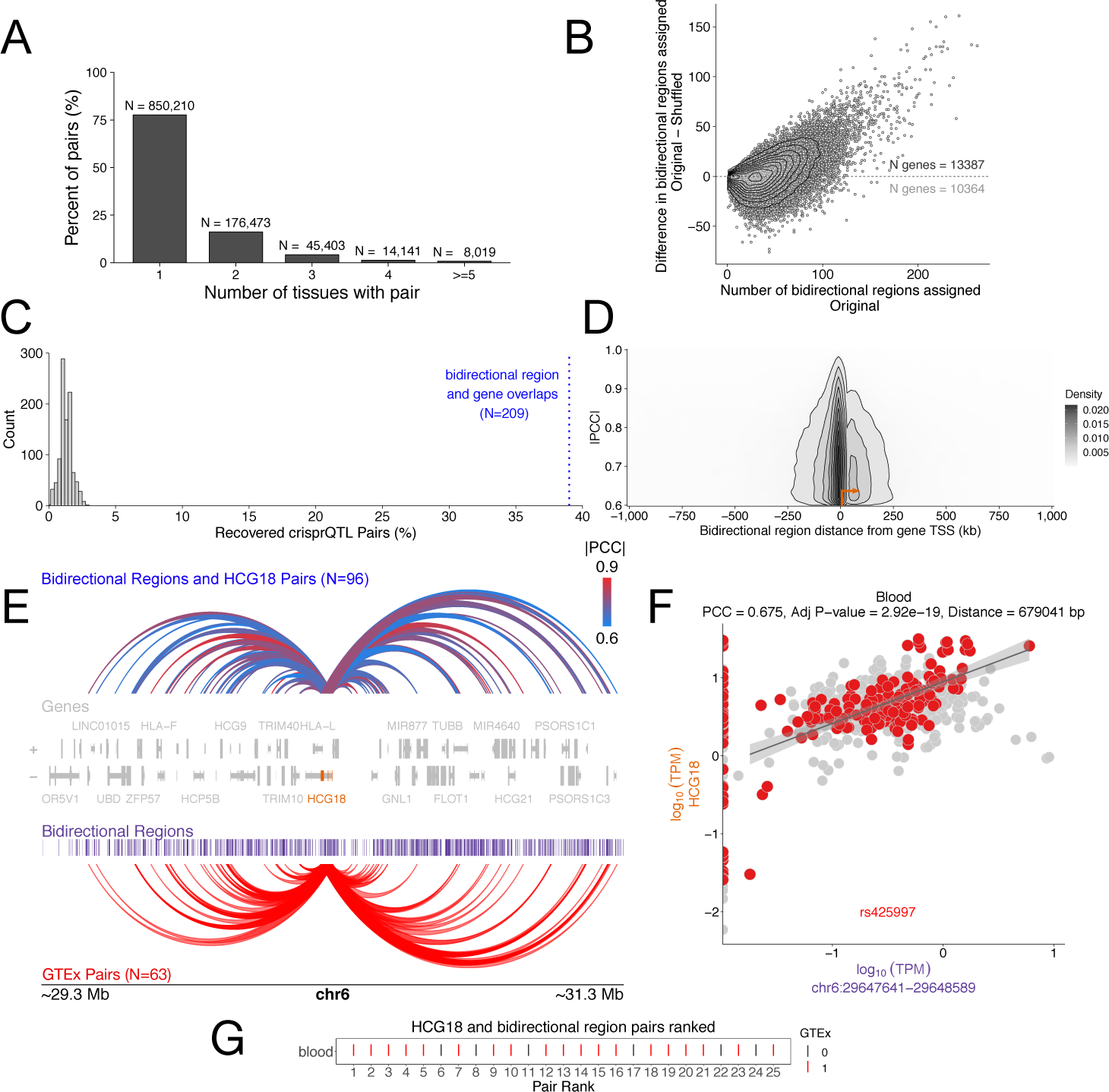
Linking bidirectional regions to protein coding genes. (A) Summary of pairs identified by tissue type (Number of unique pairs = 1,094,246). (B) Number of bidirectional regions assigned to each gene (original, x-axis) compared to the difference in number of regions assigned when shuffling (yaxis). Shuffled refers to genes correlated with bidirectional regions sampled from other chromosomes. (C) Comparison of recovered known enhancer to gene pairs[55] (blue) to pairs recovered in the shuffled approach (grey). (D) Typical distance between annotated gene and assigned bidirectional region (x-axis) and Pearson’s correlation coefficients (y-axis) for pairs overlapping significant GTEx pairs (Number of overlapping pairs = 81,046). (E) Interactions for HCG18 (NR 024052.2) with top showing identified pairs colored by absolute Pearsons correlation coefficient and bottom showing GTEx pairs. (F) Scatter plot of the raw data for one interacting pair: HCG18 (y-axis) and the bidirectional region at chr6:29,647,641-29,648,589. Samples in blood are highlighted in red. (G) Relative ranking of linked bidirectional regions on their ability to predict HCG18 levels in blood. Interactions found in GTEx are colored red.

Next, we sought to evaluate our recovered pairs by comparison to collections of known enhancer to gene linkages. First, we examined the overlap of nascent derived pairs to experimentally validated enhancer and gene pairs from K562 cells, and observed a significant recovery of known interactions (Figure 4C) [55]. Next, we considered GTEx identified pairs and found that over 80,000 nascent derived pairs overlapped with GTEx pairs (~15.28% of eQTLs pairs for variants that overlapped intergenic bidirectional regions compared to only 0.72% for random pairs), with most of these pairs near the gene TSS regions (Figure 4D). In all cases, we recover both previously identified pairs and new regions of high correlation (Supplementary Figure 24). For example, HCG18 is a long non-coding RNA that shows a high number of interactions (96 HCG18 and bidirectional pairs within 1MB of the TSS), and 65.62% of these pairs also overlap eQTLs from GTEx (Figure 4E and F)[38]. Overall, comparisons to known enhancer and gene pairs suggest that most of our identified pairs are supported by orthogonal methods.

As a final validation of our enhancer to gene linkages, we next sought to rank the bidirectional regions based on their ability to predict the transcription level of the associated target gene. To this end, we used a tree-based variable selection method[56], where the bidirectional regions were ranked based on their ability to predict the transcription levels of the paired gene. Overall, pairs found in GTEx were highly ranked across tissues (over 70% of pairs overlapping GTEx were in the top 10 of the ranks in at least one tissue) (Figure 4G and Supplementary Figure 25A). However, depending on the tissue a pair is found in, some top-ranking pairs do not overlap GTEx pairs since not all tissues in DBNascent are also in GTEx (Supplementary Figure 25B). These rankings add a separate score for each bidirectional linked to a gene in each tissue, giving us another layer of confidence in our defined pairs.

Finally, we reasoned that one use of the enhancer to gene pairs is to link intergenic disease-associated variation, typically single nucleotide variations (SNPs) to the relevant gene target. As a test of this scenario, we examined leukemia-associated SNPs from the European GWAS catalog[57]. This collection contains 7,805 distinct rsIDs (3,245 SNPs in genes and 4,560 intergenic SNPs) associated with leukemia from 35 distinct publications. When the 4,560 intergenic SNPs are assigned to the closest gene, Enricher fails to identify leukemia as a relevant term in any of the categories not built on closest gene.

In contrast to closest gene, we sought to evaluate the intergenic SNPs with our enhancer to gene pairs. Of the 4,560 intergenic SNPs, 391 were in bidirectional regions used in our correlation analysis. Of the 11 tissues, blood had the highest number of SNPs overlapping enhancer – gene pairs (Supplementary Table 4), consistent with leukemia being cancer of the blood. In blood, 126 intergenic SNPs overlapped 111 bidirectional regions, which were linked with 259 genes. Moreover, in blood – but not other tissues – Enricher showed an enrichment for leukemia-related genes in multiple categories. Finally, forty-three of the genes linked by our method also had at least one intergenic leukemia SNP, further supporting the quality of the enhancer to gene pairs. Thus, the enhancer to gene linkages inferred here were able to identify leukemia as the relevant disease, even when the closest gene approach could not.

## Discussion

Here we present DBNascent, an atlas that catalogs published nascent RNA sequencing data, with an emphasis on run-on assays. Previous work indicates that the detection of enhancer RNAs can vary by run-on protocols[58], thus we merge data across a large collection of high quality data from multiple protocols to identify sites of bidirectional transcription genome-wide across experiments, cell types, and tissues. As expected, sites of bidirectional transcription were randomly distributed across the genome, lowly transcribed, and highly variable. While previous work reports that enhancer associated transcripts are cell type specific[26, 41], our work further extends this conclusion showing they are also more specific than both protein coding genes and long noncoding RNAs. Finally, we assign cis-regulatory regions to likely target genes using a correlation based framework, identifying many possible enhancer to gene linkages.

Several previous papers showed correlations between enhancer and target gene transcription levels[18, 25–27]. Here we leveraged this fact to assign cis-regulatory regions to likely target genes using a correlation based framework. We identified correlated bidirectional and gene transcripts across human tissues. The correlated pairs we identified overlap experimentally validated enhancer—gene pairs as well as eQTLs from GTEx, supporting the use of these data to investigate regulatory region assignment. Given that the method we use to identify pairs relies on correlations of transcription levels, spurious correlations are a real concern. To curb the false positive rate, we added constraints for assigning bidirectional regions to a gene. Namely, allowed correlations that were supported by the majority of samples, pairs that were within a 1 Mb window, and had a false discovery rate of less than 0.01 on the correlation p-values. These filter steps likely enrich for true correlations but may do so at some cost with respect to less frequent but real interactions.

Additionally, we estimated the relative enrichment of true correlations relative to spurious ones by using a randomization strategy. However, it is well worth noting that there may be real correlations in the random data, as we do not control for upstream regulators (e.g. transcription factors). Despite this, we obtain more high quality correlations within the biological data and many of these pairings are good predictors of gene activity. It is well known that many disease associated variants occur in noncoding regions of the genome, often in enhancers that are associated with regions of bidirectional transcription [4, 59, 60]. Further, we demonstrate that our enhancer to gene linkages enrich for disease relevant genes better than the closest gene strategy that is commonly used to assess disease associated SNPs. Thus we recover novel biologically relevant pairings in a tissue specific manner.

Finally, it is also worth noting that the correlation linkages identified here could be used more generally to infer gene regulatory networks (GRNs). Correlation based network inference methods [61] for GRNs are, in theory, an excellent starting point for these analysis. However, our experience indicates that the increase in data set size that arises from including many potential regulatory regions makes the practical utilization of these tools challenging. Further work on building networks from these data would offer a great condition-specific resource for further experimental validation.

## Methods and Materials

All the code and methods used in the meta-analysis of nascent RNA sequencing experiments can be found on this GitHub page https://github.com/Dowell-Lab/ DBNascent Analysis. The samples were processed on a compute cluster running CentOS Linux v7.

Mouse samples were mapped to the mm10 reference genome and human samples to the hg38 genome. Databases used for comparisons that were mapped to older reference genomes were lifted to the specified genomes above using liftOver [62]. NCBI RefSeq annotations were used for both human (hg38 release GCF 000001405.40-RS 2023 03) and mouse (mm10 release GCF 000001635.26 GRCm38.p6) data.

### Nascent RNA sequencing experiments metadata collection

Nascent RNA sequencing experiments were manually curated from the Gene Expression Omnibus (GEO) [28, 29] and the Sequence Read Archive (SRA) [30]. All treatment condition times were annotated in reference to the time of cell harvest. Papers that had other high throughput experiments (including RNA-seq, ATAC-seq, ChIP-seq and 3D chromatin assays such as HiC) that were performed along with nascent RNA sequencing were noted. Two rounds of data curation were implemented where the first round was meant for data entry, and the second round for entry verification. In total, 3,638 raw samples were manually curated from 320 SRA projects (SRPs) and 287 papers. Types of data curated is described in Supplementary Table 1. Full list of papers is provided in the Supplemental References.

### Preprocessing nascent RNA sequencing experiments

All SRR accessions were downloaded from the SRA and extracted with SRA Toolkit (versions 2.8.0 and 2.9.2). Replicate information was used, where available, to combine technical replicates by concatenating fastq files. New samples resulting from technical replicate combination within a given experiment were given SRZ designations with a number equivalent to the first numerical SRR contained within. In one case, technical replicates were combined using data from multiple papers ([63, 64]) as a result of further resequencing of previously published samples. Combined samples in this case were given SRM designations, with numeric conventions the same as the SRZs. In total, 2,880 samples were generated from the original 3,638 SRR entries after technical replicate concatenation. This collection of 2,880 samples was then the source of all downstream analysis.

### Mapping reads to reference genomes

All samples in the database were trimmed, mapped to the corresponding reference genome, and assessed for sample quality using an in-house NextFlow pipeline (https://github.com/Dowell-Lab/Nascent-Flow), run with NextFlow (version 20.07.1) [65]. Briefly, within this pipeline fastq files were trimmed for adapter sequences and low quality bases using BBMap (version 38.05), then aligned to reference genomes with HISAT2 (version 2.1.0) [66, 67]. Downstream mapped read files (CRAM files and IGV-compatible TDF files) were generated with Samtools (versions 1.8 and 1.10), Bedtools (versions 2.25.0 and 2.28.0), and IGVtools (version 2.3.75), and in some cases were done so through an additional NextFlow pipeline (https://github.com/Dowell-Lab/ Downfile pipeline) [68–70]. As data was processed over an extended time frame, some software was updated resulting in changes to software versions. As such, all versions used to process a specific sample are linked to that sample in the database.

### Quality control and quality tiers

Samples were assessed for quality using metrics from the following software packages: FastQC (version 0.11.8), HISAT2 (version 2.1.0), Preseq (version 2.0.3), RSeQC (version 3.0.0), Picard tools (version 2.6.0), and BBMap[71–74]. As with mapping, specific software versions are linked to samples within the database.

Three specific metrics were used to classify samples into quality ‘tiers’ for filtering purposes: read depth after trimming, proportion of duplicates (as assessed using Picard tools), and complexity (as assessed using the modeled value for unique reads in 10 million output by Preseq). Thresholds were determined to classify samples into one of five tiers (see Figure 1), and analysis was performed on samples within tiers 1-3 unless specified otherwise.

A ‘run-on score’ was also assigned to human and mouse samples based on the following metrics: Exon/intron ratio, as calculated from the RSeQC output values for ‘CDS Exons’ and ‘Introns’, and when available from Tfit data, the total GC content of all bidirectional regions called by Tfit.

### Identifying bidirectional transcripts

Regions of nascent transcription were identified using Tfit [11] and dREG [4, 20]. Identification of regions of transcription with dREG followed the recommended pipeline (per dREG github) where uniquely mapped reads were used to generate BigWig files. BigWig input files for dREG were generated by converting filtered BAM files using bedtools bamToBed, then BED files were converted to bedGraph format using bedtools genomecov, and finally the BigWig files were generated using bedGraphToBigWig (from https://www.encodeproject.org/software/bedgraphtobigwig/) [75]. Since dREG is compute-intensive, only high-quality data sets (QC *<* 4) were processed using dREG. Identification of bidirectional transcription with Tfit followed a pipeline where multimapped reads and reads with low map quality score were filtered as shown:

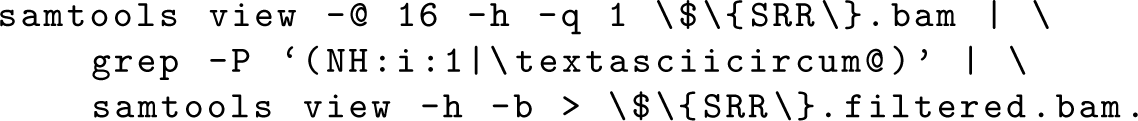

Input bedGraph files were generated using genomeCoverageBed from bedtools. Tfit was run in a two step processes, first with the template matching module to identify sites of bidirectional transcription, then these regions were used as input to fit the precise RNA polymerase behavior. The nextflow pipeline used for characterizing bidirectional transcription with both Tfit and dREG can be found on GitHub (https://github.com/Dowell-Lab/Bidirectional-Flow).

### Merging regions of bidirectional transcription

#### Updated muMerge method

This project required the development of several new features for *muMerge*[19] in order to facilitate aspects of the analysis. Namely, the creation of a filter to remove regions occurring in only one sample (“singletons filter”; -r, --remove singletons) and an option to save a record of which samples contribute to an individual mumerged region (“save sample IDs”; -s, --save sampids). The singletons filter filters any output region supported by one one data input. The save sample IDs option adds a fourth column to the output file which contains a comma separated list of all the sample IDs that contributed at least one loci to the calculation of that given mumerged region. These sample IDs are reported in alphanumeric order. These two features have been included in mumerge v1.1.0 (see https://pypi.org/project/mumerge/ for details).

#### muMerging samples across conditions

Regions (from replicates, conditions, and bidirectional calling methods) were merged using *muMerge* (described above). Since *muMerge* combines regions in a probabilistic way, replicate information and sample conditions were taken into account for the merging processes. Tfit and dREG bidirectional transcript calls were first mumerged separately by paper based on the experimental setup (that is by cell/tissue type, experimental condition and replicate information).

The *muMerge* bed files by experiment/paper were combined based on the cell/tissue types for Tfit and dREG, where the same cell/tissue types were treated as “replicates” and the different cell/tissue types were the “conditions”. All samples were mumerged and calls were filtered based on the paper QC and the GC content of the 300bp region around the center of the call (Supplemental Figure 5). Bidirectional calls from papers with GC content ¿ 0.49, average paper QC score ¡ 3 were kept (Supplemental Figure 5). In summary, samples from 61 mouse papers and 101 human papers were used for the final species-specific *muMerge* steps. Furthermore, the dREG and Tfit *muMerge* files were combined such that Tfit calls were used for overlapping regions creating a master *muMerge* file for both species (shown schematically in Figure 2A). Lastly, the dREG and Tfit master *muMerge* file was filtered. Regions that were greater than 150bp and less than 2.5kb were kept in the human bidirectional calls (less than 2kb in mouse regions). The minimum read length of 150bp was selected as this is the maximum read length for libraries in the nascent RNA samples in the database, and the upper limit was selected so that the largest regions from dREG and Tfit matched. In order to remove false positive bidirectional calls, bidirectional calls that were in regions of converging gene transcripts (where converging genes here are defined as sense and antisense transcripts that collide) within 1kb were also removed. Some general observations on the two methods were made. First, we found that dREG broke bidirectional regions identified by Tfit into smaller chunks. Despite this,

Tfit calls slightly more regions per sample compared to dREG (Supplemental Figure 4A). Second, both methods struggled to call bidirectional regions within introns when the gene was transcribed robustly, though generally dREG called more in these regions (Supplemental Figure 26). Finally, when samples have poor quality, both dREG and Tfit tend to call a higher percent of gene TSS regions as these are more highly transcribed and therefore have the most robust signal (Supplemental Figure 27). More generally, we see a higher number of TSS regions called by dREG comapred to Tfit. As no gold standard exists for transcribed regulatory elements, it is unclear which of the two methods is more accurate. There are regions that are recovered by both methods, and regions unique to each bidirectional transcript caller (Supplemental Figures 4B and C). So in this paper we combined calls from both methods, keeping track of their origin, i.e. bidirectional caller.

The processing pipeline was performed with R (version 3.6.0), using the package data.table (version 1.14.2), and genome arithmetics were performed using bedtools [76, 77]. The pipeline used can be found on GitHub here: https://github.com/Dowell-Lab/ bidirectionals merged.

### Summary of transcription

#### Bidirectional transcripts overlapping genomic features

Annotated bidirectional transcripts were overlapped with RefSeq genome feature using bedtools. The bidirectional coordinates were overlapped with intron, promoter (1kb upstream of a gene TSS), exon and intergenic regions (regions not annotated in the reference annotations). Across all features, the minimum fractions overlap required per region was 0.5 as shown:

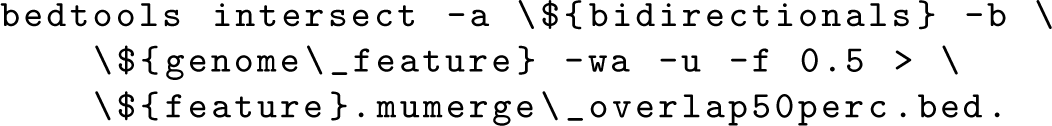

The percent of overlap in each feature category was calculated as a fraction of the total bidirectional transcripts (847,521 in human and 680,735 in mouse).

#### Gene summary statistics methods

All gene statistic analyses were performed using R (version 4.3.0), the R package data.table (version 1.14.8), and bedtools (version 2.30.0).

To identify TSS bidirectional regions we used bedtools intersect to find overlaps of the final bidirectional calls with a 600bp window around the TSS (300bp both directions). For gene transcripts with multiple bidirectional regions overlapping, the bidirectional whose center (*µ*) was closest to the TSS is considered the TSS bidirectional. The notebook corresponding to this analysis can be found on GitHub here: https://github.com/Dowell-Lab/bidirectionals merged/notebooks/.

To further interrogate the number of intronic and exonic bidirectional regions at a gene-centric level, we used bedtools to intersect a 100bp window around *µ* with introns and exons, requiring that the bidirectional have majority coverage in the exon or intron. The following code was used:

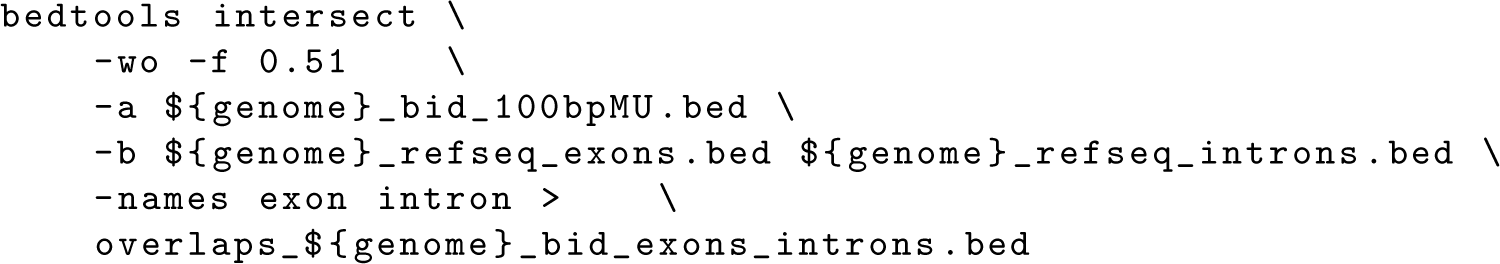

To get percentile based coordinates of bidirectional regions within genes, all positions (midpoint, start, and end of bidirectional and gene) were transformed in relation to the gene itself, with the TSS marking 0 and PAS marking the length of the gene. These coordinates were then multiplied by a size factor calculated as shown below to standardize the coordinates as percentiles where *i* refers to the transcript isoform.

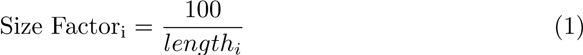

where

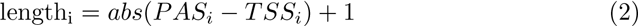

The notebook corresponding to this percentile analysis can be found on GitHub here: https://github.com/Dowell-Lab/DBNascent Analysis/analysis.

### Overlapping bidirectional transcripts with candidate cis-regulatory elements

#### Region overlaps

Regions of bidirectional transcription were overlapped with candidate cis-regulatory element (cCRE) databases (ENCODE, EnhancerAtlas, FANTOM5 and eQTL data) using bedtools [3, 32, 33, 37]. The reprocessed version 9 of FANTOM5 data for mm10 and hg38 were downloaded from the FANTOM website https://fantom.gsc.riken.jp/5/ datafiles/reprocessed/. EnhancerAtlas 2.0 data was download from the database website http://www.enhanceratlas.org/data/download/species enh bed.tar.gz and using liftOver, coordinates were converted to the mm10 or hg38 for mouse or human respectively. ENCODE candidate cis-regulatory elements were downloaded from the UCSC genome browser for both human http://hgdownload.soe.ucsc.edu/gbdb/hg38/ encode3/ccre/encodeCcreCombined.bb and mouse https://hgdownload.soe.ucsc.edu/ gbdb/mm10/encode3/ccre/encodeCcreCombined.bb. Finally, significant eQTLs from GTEx version 8 (GTEx Analysis v8 eQTL.tar) were downloaded from the GTEx portal (https://gtexportal.org/home/). Mouse eQTLs were downloaded from the from Gonzales et al. paper [78].

Regions of overlap were calculated using the minimum overlap of 1bp as shown:

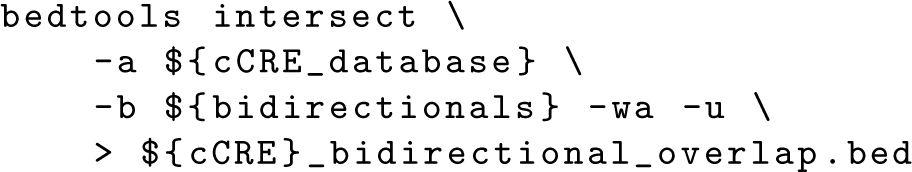

The percent overlap was calculated with respect to the database size.

#### Odds ratio with GTEx

Bidirectional transcription regions were overlapped with disease associated variants from GTEx (version 8) and odds ratio calculated [38]. All eQTLs including non-significant instances (GTEx Analysis v8 eQTL all associations) were downloaded from the google cloud location stated in the GTEx portal. The odds ratio was calculated by counting the number of GTEx eQTL variants that overlapped bidirectional transcripts for both significant and non-significant variants and getting the fraction of the variant overlapping bidirectional transcripts versus not overlapping the transcripts (Supplement Figure 13).

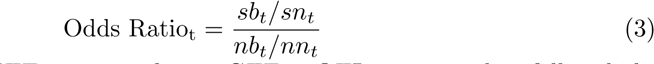

Where *t* is a given GTEx tissue, *sb_t_* are GTEx eQTL variants that fall in bidirectional transcripts, *sn_t_* are significant variants that fall outside of bidirectional transcripts, *nb_t_* are non-significant variants that fall within bidirectional regions and *nn_t_* are non-significant GTEx variants that do not fall in bidirectional regions. The odds ratio calculation was performed using the library epitools (version 0.5-10.1) in R [79].

#### Calculating base content

Base composition for bidirectional transcript calls from dREG and Tfit was defined as the ratio of GC content in the center 300bp (typically contains the RNA loading position) relative to larger bounding 3kb region. This was performed by extracting sequences using bedtools getfasta, and counting the frequency of bases (A/T/C/G) within the small (300bp) and large (3000bp) regions. This counting was performed in python 3.6 [80].

#### Counting reads

Reads were counted using featureCounts from RSubread (version 2.12.3) [81]. For gene transcripts, sense strand reads over gene bodies were counted as these show more consistency across nuclear run-on protocols [58]. Gene bodies were defined for genes over 300bp long as full gene lengths from the TSS to the TES, truncated at the 5*^1^* end by 30% of the gene length up to a maximum of 750bp truncation. Genes

*<* 300bp were not truncated. For bidirectional regions, reads on both strands were counted across the entire called region using the coordinates start+1 to end-1. In both cases, multimapping reads were ignored. Multi-feature overlap was allowed for counting across genes but not bidirectional regions.

#### Normalizing read counts

Normalization of counts was done using transcripts per million (TPM) normalization as shown below [82, 83]:

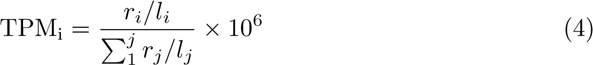

where *r_i_* are the mapped reads for transcript *i* (for all genes and bidirectional transcripts), *l_i_* is the transcript length and Σ_*j*_*r*_*j*_/*l*_*i*_ sums all *j* length normalized transcripts. The ratio is multiplied by a scaling factor of 10^6^. The counts for genes was normalized over full length of the longest transcript and the 5*^1^* end of that transcripts were truncated. To avoid double counting reads, for bidirectional transcripts that overlap genes, the opposite sense read counts were used in the normalization step. In summary, the total number of transcripts included 5*^1^* end truncated genes, intergenic bidirectional transcripts and intragenic bidirectional regions where counts on the opposite strand of gene were used for these bidirectional transcripts.

#### Calculating summary statistics

The summary statistics described below were calculated using R. For all samples, the average and median transcription values were calculated based on the normalized counts.

#### Coefficient of variation

Across-sample coefficient of variation (CV) for human samples were calculated as follows:

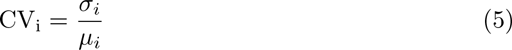

where for transcript *i* the standard deviation (*σ_i_*) for normalized counts is divided by the average normalized counts (*µ_i_*).

#### Principle component analysis

The principle component analysis (PCA) was performed using human GRO-seq and PRO-seq samples with QC 1 and 2 and with a GC content greater than 0.49. This resulted in 751 samples for analysis. RefSeq hg38 genes with high CV (greater than 3) were used for the PCA. Since bidirectional transcripts are lowly transcribed and have a higher coefficient of variation, more stringent filter was used to filter these transcripts (CV greater than 6 and average TPMs greater than 0.1). In both cases, normalized counts were log transformed, euclidean distances calculated using the R package distances (version 0.1.8), and PCA performed with the prcomp function in the stats package in R [84].

#### Tissue specificity

For analysis of variation and tissue specificity, genes were classified into ‘coding’ and ‘noncoding’ genes according to ‘NM ’ vs ‘NR ’ accession number prefixes. Bidirectional regions were classified into ‘promoter’, ‘exonic’, ‘intronic’, and ‘intergenic’ bidirectional regions according to bedtools overlaps with those annotations as described above, with exonic, intronic, and intergenic bidirectional regions also being lumped together as ‘nonpromoter’ bidirectional regions for some comparisons.

SPECS scores were calculated using a custom python-based implementation of the method described previously [40], using Python (version 3.6.3), Numpy (version 1.19.2), Pandas (version 1.0.2), and Scipy (version 1.4.1). This script can be found in the GitHub repository https://github.com/Dowell-Lab/DBNascent Analysis/ [85–88].

#### Correlation and Co-transcription Analysis

Building the co-transcription bidirectional and gene pairs from nuclear run-on data was split into three steps: (1) finding pairs of highly correlated genes and bidirectional transcripts, (2) removing bidirectional transcripts that are in the downstream regions of genes and (3) filtering high confidence pairs (See Supplementary Figure 17).

### Step 1: Pairwise correlation of gene and bidirectional transcripts

Pearson’s correlation coefficients between genes and bidirectional regions were calculated using WGCNA (version 1.70-3)[61, 89] and all the filtering done in R. Comparison were among transcribed regions within a chromosome (i.e. no cross chromosome comparisons). The input to WGCNA was normalized counts for genes where the 5*^1^* end was truncated (as described in the “Counting reads” section) along with bidirectional regions. These counts were log transformed as shown below.

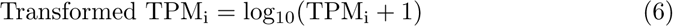

Additionally, transcripts with zero counts were excluded from the pairwise calculations. Given the samples with transcription, the Pearsons correlation coefficient (PCC) was calculated as follows:

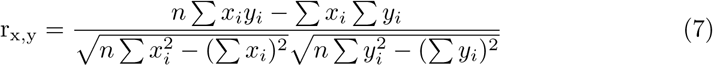

transcribed for both gene and bidirectional regions. The p-value was calculated from the Student’s t-distribution and the *t* statistic was calculated as:

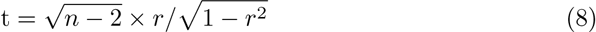

where *n* are the number of samples and *r* is the PCC. The output from the correlation calculations included the transcript identifiers, distance between the center of the bidrectional and the TSS and TES of a gene, the PCC, the p-value and adjusted p-values [90]. Additionally, the number of samples used in the correlation calculation as well as the tissue a pair is identified in are reported.

The code used to calculate pairwise correlations can be found on this GitHub repository https://github.com/Dowell-Lab/bidir gene pairs.

### Step 2: Filtering bidirectional transcripts

It has been shown that gene transcription from nuclear run on assays shows transcription past the polyadenylation side (PAS) [2, 46]. Since bidirectional transcripts in these regions would contain reads from the gene, it is difficult to disentangle signal of the bidirectional transcript from the gene transcripts. This would result in high correlation between these bidirectional regions and the gene upstream. Therefore, pairs with bidirectional transcripts that were with 15kb downstream of the PAS were removed.

### Step 3: Filtering for high confidence pairs

To ensure robustness of the correlations, we filtered our tissues to those that include at least 15 samples, leaving 11 tissues to assess. In each tissue, significant bidirectional and gene pairs were defined as pairs with 1Mb (from the center of the bidirectional transcript to the gene TSS), where the absolute Pearsons correlation coefficient (PCC) was greater than 0.6, and the adjusted p-value was less than 0.01 (Supplementary Figure 17). Importantly, pairs had to be supported by over 10 samples, in order to curb spurious correlations.

#### Evaluation of relative false positive rate

Since it has been shown that most enhancers regulate gene expression in a distance-dependent manner and within topologically associated domains (TADs) [91, 92], we reasoned that linking genes with bidirectional transcripts from outside TADs would yield an estimation of false positive links. Thus, we reasoned that genes and bidirectionals on distinct chromosomes would be highly unlikely to be real pairs. Using this, we assessed the relative false positive rate using a randomization strategy. Specifically, when considering a gene on a given chromosome, we consider all bidirectionals from the same chromosome (e.g. relative distance to a bidirectional region is unaltered) but sampled a random transcription vector from the set of all bidirectionals not on the chromosome. For each chromosome, the shuffling of bidirectional regions from the remaining chromosomes was done without replacement. The shuffling process was done in R using the sample function in the base package. The correlation analysis and filtering of pairs was done with the same methods described above (See correlation methods). The code used for the shuffling method can be found on GitHub (https://github.com/Dowell-Lab/DBNascent Analysis).

#### Overlap of pairs with eQTLs and crisprQTLs

Gene and bidirectional pairs derived from the co-expression method were overlapped with pairs from crisprQTLs validated enhancer – gene pairs from Gasperini and company [55]. The gene and bidirectional pairs were randomly shuffled 1000 times within each chromosome, and the overlaps with the crisprQTLs assessed. The percent overlap was calculated based on crisprQTLs that were present and transcribed in our dataset (536 out of the 664 tested crisprQTLs). The random pairs and true pair overlaps were compared and plotted as a histogram (See Figure 4C). For eQTLs from GTEx, the randomization was only done once [38]. The eQTLs/crisprQTLs were overlapped with bidirectional transcript coordinates using bedtools intersect and the genes were matched based on the gene name in R. The shuffling was done using the sample function from the base package in R.

#### Ranking bidirectional transcripts with GENIE3

To rank bidirectional transcripts assigned to each gene in each tissue, the normalized counts for bidirectional regions and their paired gene were used as input to the GENIE3 (version 1.22.0) library in R (version 4.3.1) [56]. The bidirectional transcripts were labeled as regulators. Regression trees were learned using random fores ts, and the *p* is the number of bidirectional transcripts (i.e. regulators). The number of trees that were grown per ensemble was 1000. The output from this analysis returned a list of gene and bidirectional pairs in each tissue, the rank for each bidirectional transcript, and the weight of link.

#### GWAS SNPs connected with correlation pairs

We used custom python software with the pandas (version 2.1.3) library to link SNPs to genes. GWAS-detected leukemia SNPs were downloaded from the NHGRIEBI GWAS Catalog (EFO 0000565) [57]. SNPs (rsIDs) were filtered to remove those without position and deduplicated (12,629 total rsIDs and 7,805 rsIDs after deduplication). Bedtools was used to find the closest gene to each SNP and to determine which SNPs were intergenic. Bedtools was also used to determine which SNPs were within a bidirectional region in DBNascent. SNPs found in intergenic bidirectional regions were overlapped with DBNascent bidirectional region – gene pairs. When evaluating for enrichment, the gene within the pair or the closest gene was fed to Enrichr[93]. DBNascent-pair genes were loaded to Enrichr with the background of all genes identified in any pair within blood. The closest gene list showed enrichment for leukemia only in gene lists that utilized closest gene in their construction, such as PhenGenI Association, GWAS Catalog 2023, DisGeNET, and GeDiPNet[94–96]. While DBNascent-paired genes showed enrichment for leukemia in both the OMIM and OMIM enriched categories [97]. OMIM links genes to diseases via literature and OMIM-enriched adds protein-protein interactions to the literature.

The code used for this analysis can be found on GitHub here https://github.com/Dowell-Lab/snpconnecter.

### Data visualization

#### Plot generation

Plots were generated using ggplot2 (versions 3.6.0 and 3.4.2), cowplot (version 1.1.1), ComplexUpset (version 1.3.3) and VennDiagram (version 1.7.3) packages in R [98–100].

#### Genome tracks

Genome tracks were generated with pyGenomeTracks (version 3.8) in python [101, 102]. Gene and bidirectional pairs were represented using the plotgardener (version 1.6.4) package in R with hg38 gene annotations from TxDb.Hsapiens.UCSC.hg38.knownGene (version 3.17.0) [103, 104].

#### Illustrations

Illustrations were generated using the open-source tool Inkscape (version 1.0.1).

#### Nascent Database Structure : Back-end

All scripts for database construction and maintenance, as well as a visual schema of the database, can be found at https://github.com/Dowell-Lab/DBNascent-build.

The MySQL database backend for DBNascent was built using Python and SQLAlchemy (version 1.4.31). All metadata collected was stored in text files divided by paper (for experiment-level metadata) and samples within a paper (for sample-level metadata), which were then read to parse into database tables. Broader metadata detailing organism and cell type/tissue information was also manually curated into tables, available in the repository. Quality control metrics output by the software packages FastQC, HiSAT2, Picard tools, Preseq, RSeQC, and Pileup were pulled directly from files output by those packages. Software version information and bidirectional summary statistics were pulled from pipeline outputs from our Nextflow pipelines (NascentFlow and BidirectionalFlow), detailed previously.

A front-end website for DBNascent (nascent.colorado.edu) was built in Python 3.6.8 using the packages Django (version 3.2.16) and uWSGI (version 2.0.21) and is served using nginx (version 1.20.1). The website is maintained by the IT group at the BioFrontiers Institute.

### Data Availability

Processed data and intermediate files can be found on Zenodo (10.5281/zenodo.10223322) and on the DBNascent website (nascent.colorado.edu).

## Supplementary information

1. Supplemental Figures
2. Video File 1

## Funding

This work was funded by the National Science Foundation under grants ABI1759949 and the National Institutes of Health grant GM125871 and HL156475.

## Competing interests

The authors declare that they have no competing interests.

## Author’s contributions

RFS, MAA and RDD conceived and designed the analysis. RFS and LS collected data and managed metadata curation. RFS performed analysis with help from LS and ZLM to construct DBNascent, LS for cell type specificity analysis, TJ for GC analysis, MAA for linking SNPs to genes, and HAT in feature overlap characterization. JTS revised *muMerge*. RFS, LS, RDD wrote the paper. All authors revised manuscript.

## Acknowledgements

We thank Joseph Cardiello, Samuel Hunter, Jesse Kurland, Kendra Meer, Marko Melnick, Daniel Ramirez, Antonio Salcido-Alcantar, Gilson Sanchez, Jessica Westfall, Qing Yang and Chi Zhang for contributions to meta-data curation. We thank Charles Danko for assistance and advice regarding running dREG. We are also grateful to the BioFrontiers IT department for their support in building the database.

## Code availability

All the code and methods used in the meta-analysis of nascent RNA sequencing experiments can be found on this GitHub page https://github.com/Dowell-Lab/ DBNascent Analysis. All scripts for database construction and maintenance, as well as a visual schema of the database, can be found at https://github.com/Dowell-Lab/ DBNascent-build. Processed data and intermediate files can be found on Zenodo (10.5281/zenodo.10223322).

- Database backend: https://github.com/Dowell-Lab/DBNascent-build
- Data preprocessing: https://github.com/Dowell-Lab/Nascent-Flow
- Bidirectional calling and read counting: https://github.com/Dowell-Lab/ Bidirectional-Flow
- muMerge and combining bidirectional regions: https://github.com/Dowell-Lab/bidirectionals merged
- Correlation of bidirectional regions and genes: https://github.com/Dowell-Lab/ bidir gene pairs
- Downstream analysis of pairs: https://github.com/Dowell-Lab/DBNascentAnalysis
- Linking GWAS SNP to genes with DBNascent: https://github.com/Dowell-Lab/ snpconnecter

## Notes

https://nascent.colorado.edu/home/

### Competing Interest Statement

The authors have declared no competing interest.

