## Supplemental Figures for "Atlas of nascent RNA transcripts reveals enhancer to gene linkages"

December 7, 2023

- [1] BioFrontiers Institute, University of Colorado, Boulder CO 80309 USA
- [2] Department of Computer Science, University of Colorado, Boulder CO 80309 USA
- [3] Department of Molecular, Cellular and Developmental Biology, University of Colorado, Boulder CO 80309 USA

### Supplementary Note

All citations included in DBNascent are listed on the website and included as References in this file.

### Supplementary Figures

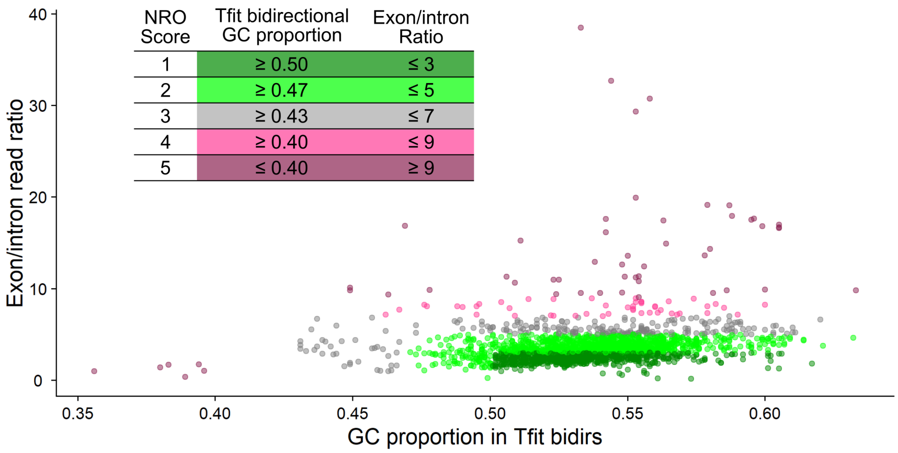


Figure 1: **Distribution of nuclear run-on (NRO) scores for human and mouse samples in DBNascent.** As nuclear run-on signals are ideally pre-splicing, the exon to intron ratio is expected to be low. Higher ratios are indicative of issues with the nascent RNA enrichment. Notably, Tfit output were not available for some very poor quality samples. In these scenarios, the NRO score was based on exon/intron ratio, calculated from RSeQC metrics.

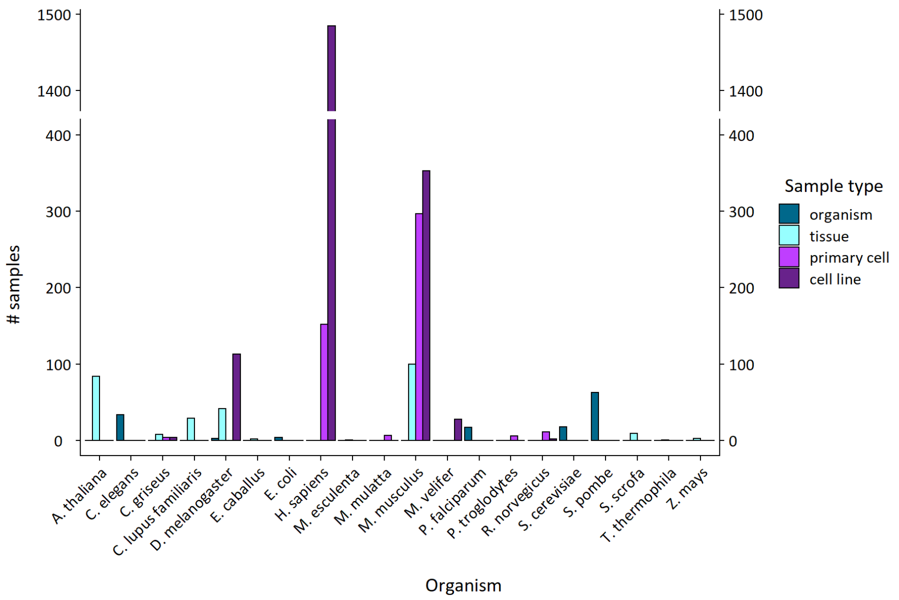


Figure 2: **Sample types represented in DBNascent.** The most represented cell type across the database are cell lines, followed by primary cells. Most of these sample types originate from human and mouse samples.

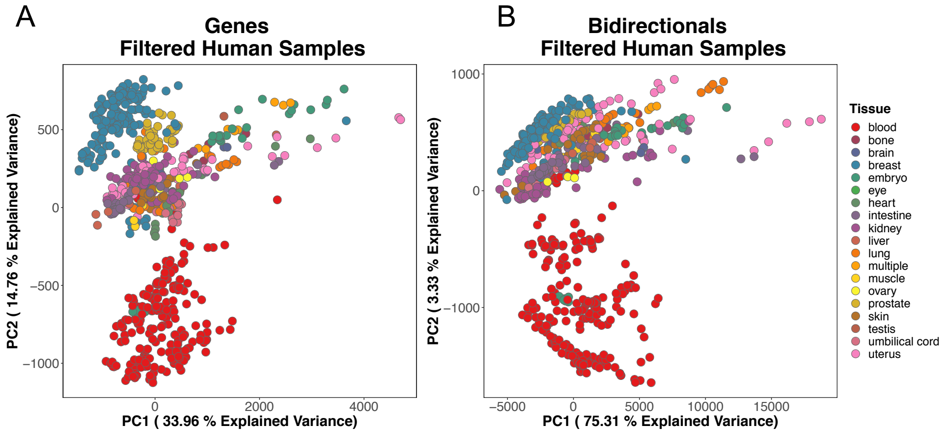


Figure 3: **PCA plot for human genes and bidirectional regions.** PC1 and PC2 for (A) RefSeq hg38 genes and (B) bidirectional regions using 751 high quality samples colored by tissue type.

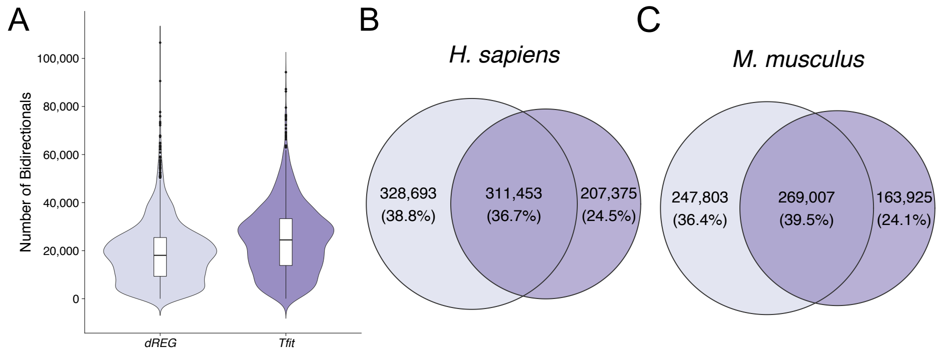


Figure 4: **Summary of bidirectional calls from dREG and Tfit.** (A) Distribution of the number of bidirectional regions called by both dREG and Tfit for each sample (number of samples in human = 1638 and mouse = 750). Overlap of (B) human and (C) mouse bidirectional calls after *muMerge*.

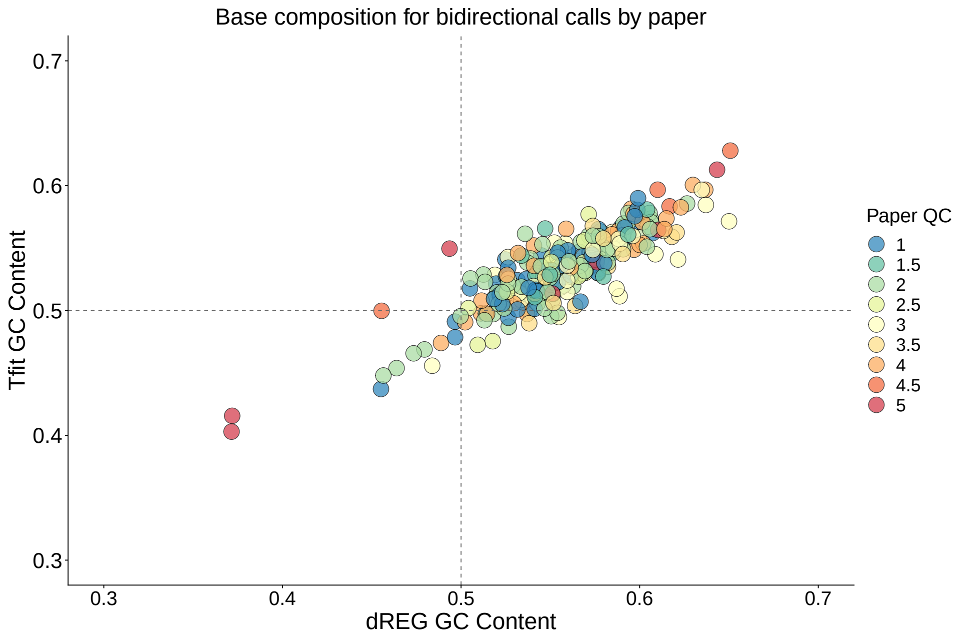


Figure 5: **Base composition for papers after *muMerge*.** Average GC content around center of called bidirectional region for each paper, by caller (x-axis: dREG, y-axis: Tfit).

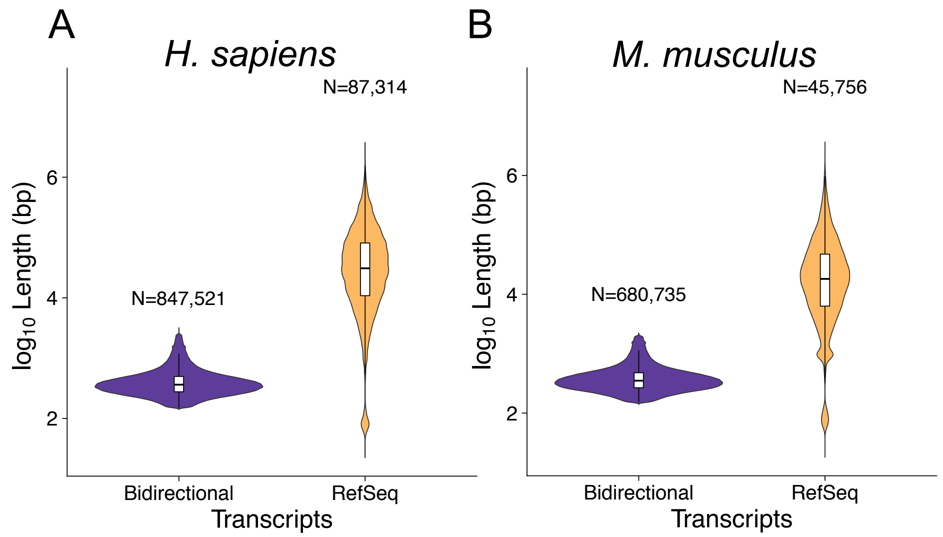


Figure 6: **RefSeq gene annotations and bidirectional transcripts length distributions.** (A) Human and (B) mouse transcript length distributions for bidirectional and RefSeq (hg38 and mm10 respectively) annotations.

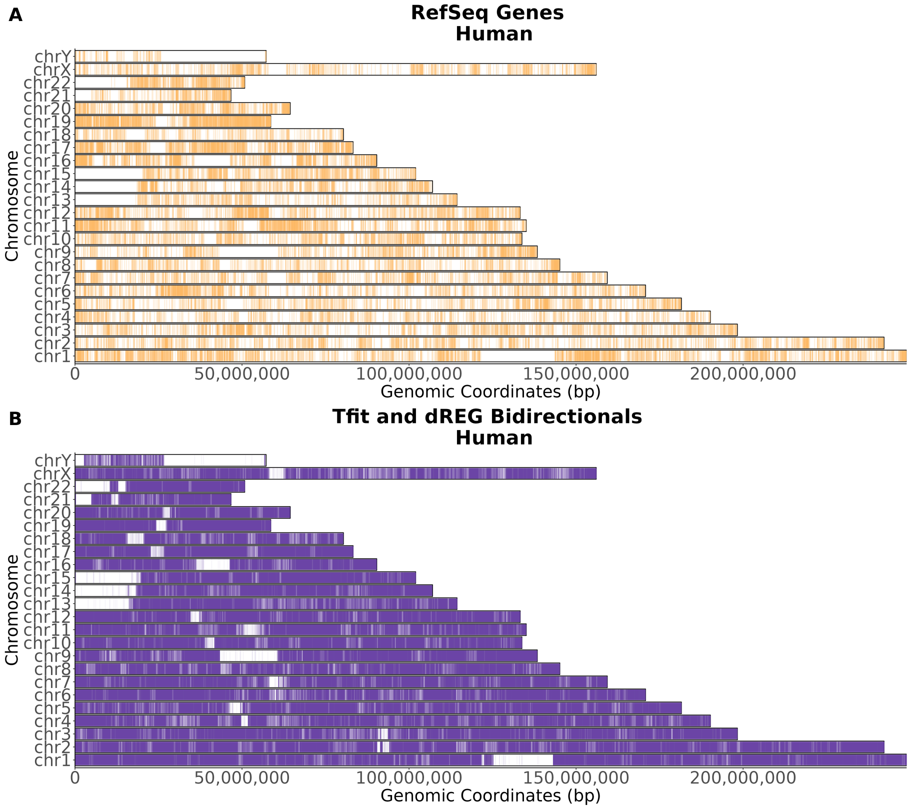


Figure 7: **Human transcribed regions.** (A) RefSeq hg38 genes and (B) bidirectional regions (N = 847,521) transcribed marked as a point on each chromosome.

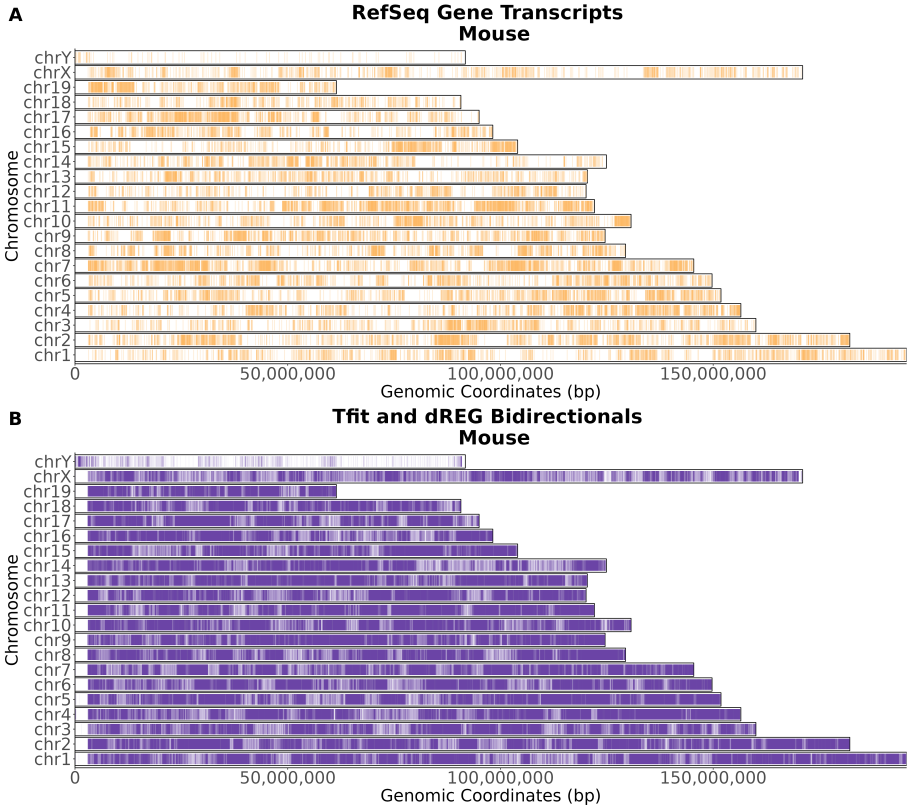


Figure 8: **Mouse transcribed regions.** (A) RefSeq mm10 genes and (B) bidirectional regions (N = 680,735) that are transcribed marked as a point on each chromosome.

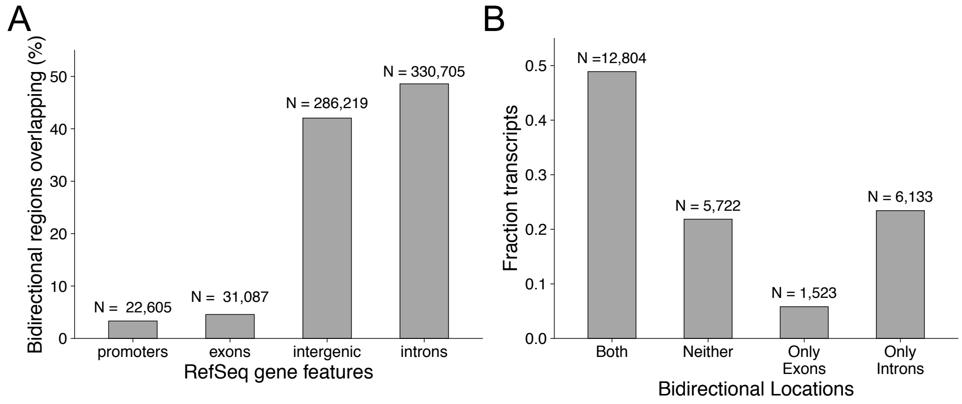


Figure 9: **Overlap between mouse bidirectional regions and RefSeq gene annotations.** Mouse overlaps similar to Figure 2B-C. (A) Percent of bidirectional regions overlapping RefSeq mm10 annotations. (B) Fraction of RefSeq mm10 gene features with bidirectional transcripts.

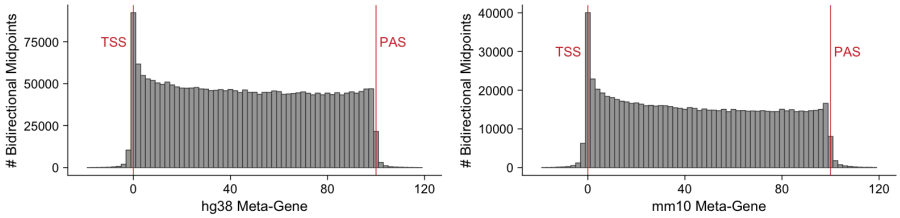


Figure 10: **Distribution of bidirectional midpoints across a gene.** Metagene plots showing midpoint position of bidirectional regions relative to genes, normalized to percentiles of gene length. Red labels highlight the annotated transcription start site (TSS-0) and polyadenylation site (PAS-100). Left is for human transcripts with an overlap (RefSeq hg38 annotations - 23,752 genes and 80,806 transcripts). Right is for mouse transcripts with an overlap (mm10 annotations - 21,590 genes and 40,652 transcripts).

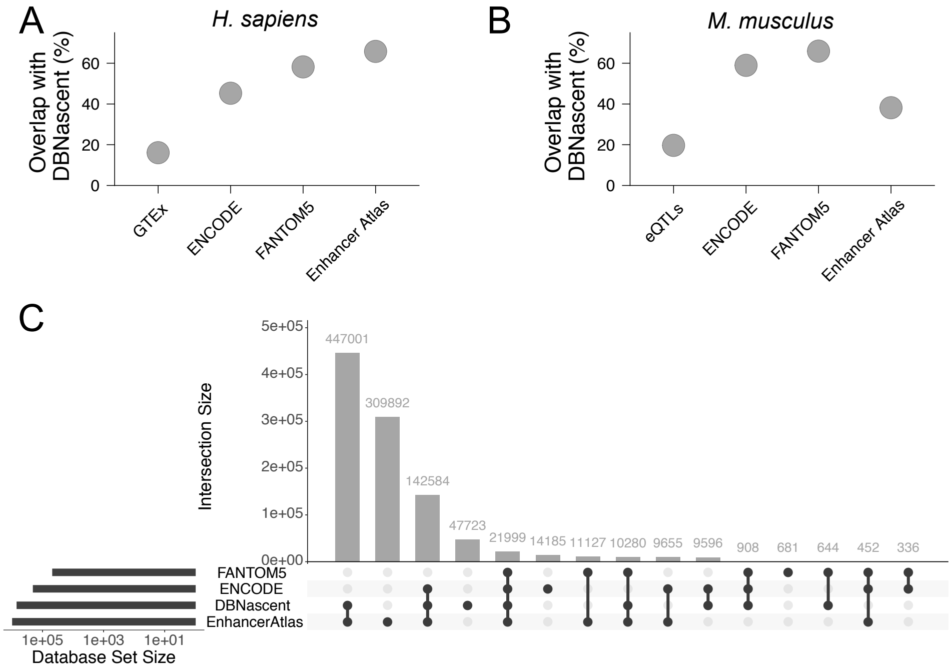


Figure 11: **Overlap between bidirectional regions and candidate enhancer databases.** (A) Human and (B) mouse overlap of other cis-regulatory region databases (eQTLs, ENCODE, FANTOM5 and Enhancer Atlas) with bidirectional regions. (C) Mouse bidirectional regions overlapping with other databases.

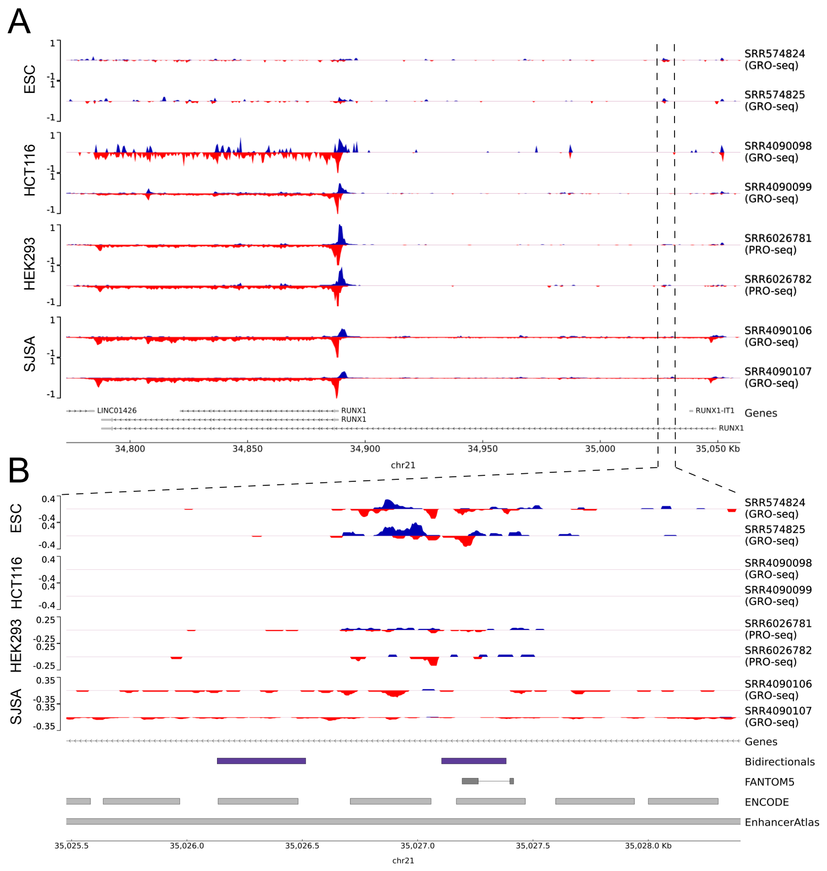


Figure 12: **RUNX1 and bidirectional region genome track.** An example of an intronic bidirectional region, in this case in RUNX1. (A) Coverage across four cell lines (ESC, HCT116, HEK293 and SJSA). Region shown chr21:34,772,887-35,059,610. (B) Zooming into the bidirectional region in the first intron for RUNX1. Red: positive strand reads, Blue: negative strand reads. Region shown chr21:35,025,477-35,028,400.

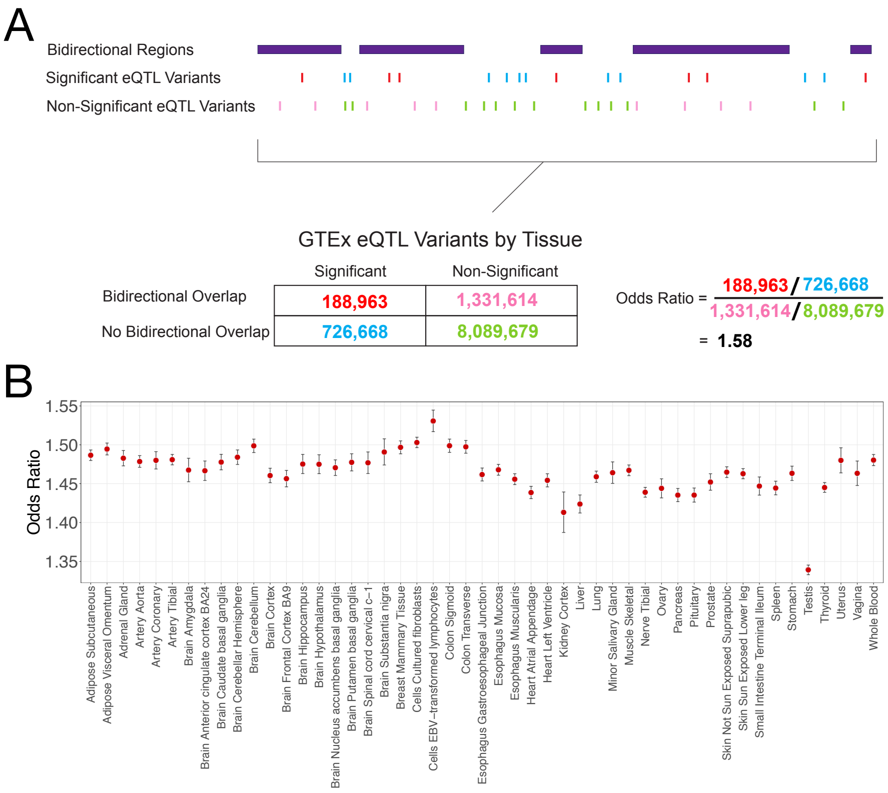


Figure 13: **Odds ratio for GTEx eQTLs and bidirectional transcripts overlapping.** (A) Schematic for odds ratio calculation. Called bidirectional regions (purple boxes) are compared to eQTL information (both significant and non-significant calls). eQTLs are color labeled by overlap (red: significant and overlap; blue: significant but no overlap; pink: non-significant and overlap; green: non-significant and no overlap). (B) Odds ratio between significant GTEx eQTL variants that overlap bidirectional regions compared to non-significant variants overlap for all tissues in GTEx.

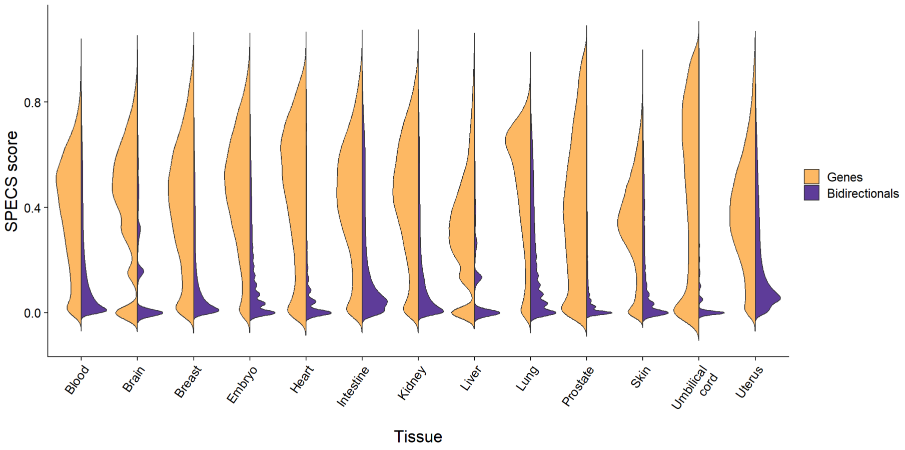


Figure 14: **Distribution of SPECS scores across genes and bidirectional regions for each tissue present in DBNascent.** Only tissues with $> 5$ total samples of QC score $< 4$ were considered.

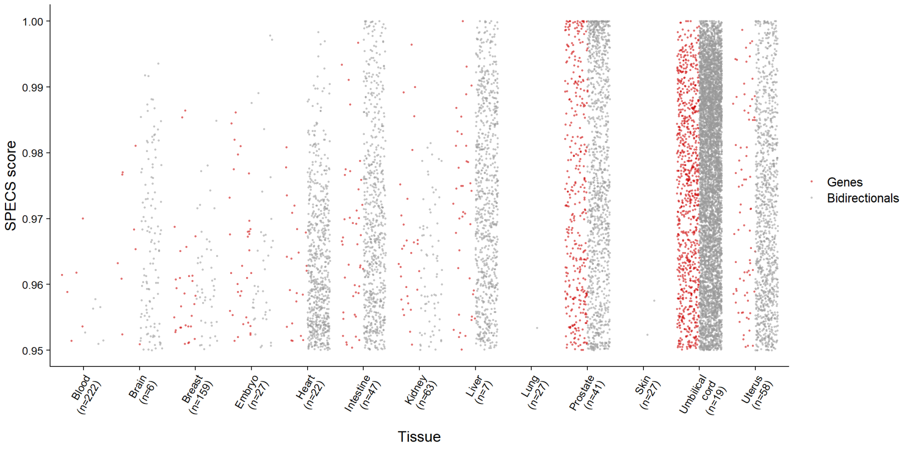


Figure 15: **Highly specific genes and bidirectional regions based on SPECS score.** The n-value corresponds to the number of samples comprising a specific tissue. A SPECS score of greater than 0.95 is considered highly specific for a tissue. In general, the relative proportions of high SPECS scores genes and bidirectional regions correlate across tissues.

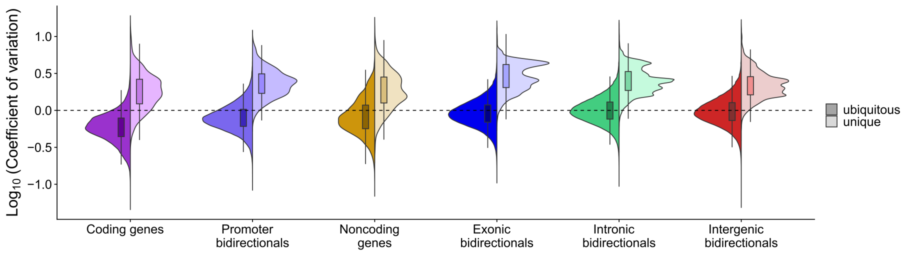


Figure 16: **Coefficient of variation across region class among ubiquitous and tissue unique regions.** Ubiquitous regions are transcribed in all 13 tissues analyzed, whereas unique regions are only transcribed in a single tissue.

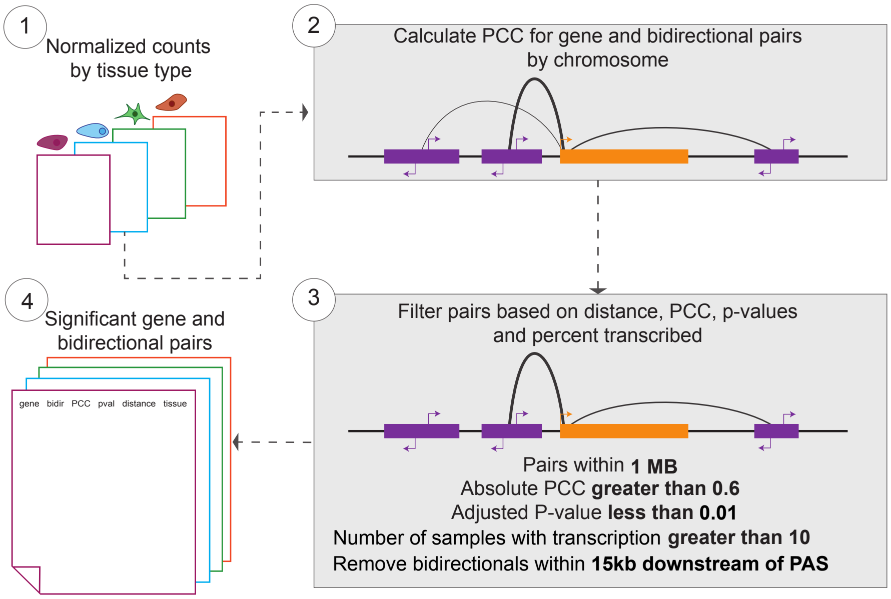


Figure 17: **Tissue specific correlations.** (1) Counts across transcripts are normalized by library size and transcript length. The samples are separated by tissue type (Blood, Breast, Embryo, Intestine, Kidney, Lung, Prostate, Skin, Umbilical cord, Uterus, Heart). (2) Within each tissue, correlations between genes and bidirectional transcripts are computed. (3) The pairs are filtered based on distance (less than 1Mb), Pearson's correlation coefficient (PCC) greater than 0.6 or less than -0.6, an adjusted p-value less than 0.01 and the pairs should be supported by 10 or more samples. Lastly, bidirectional regions with 15kb downstream on the polyadenylation site (PAS) were removed (see Methods). (4) Significant pairs are saved.

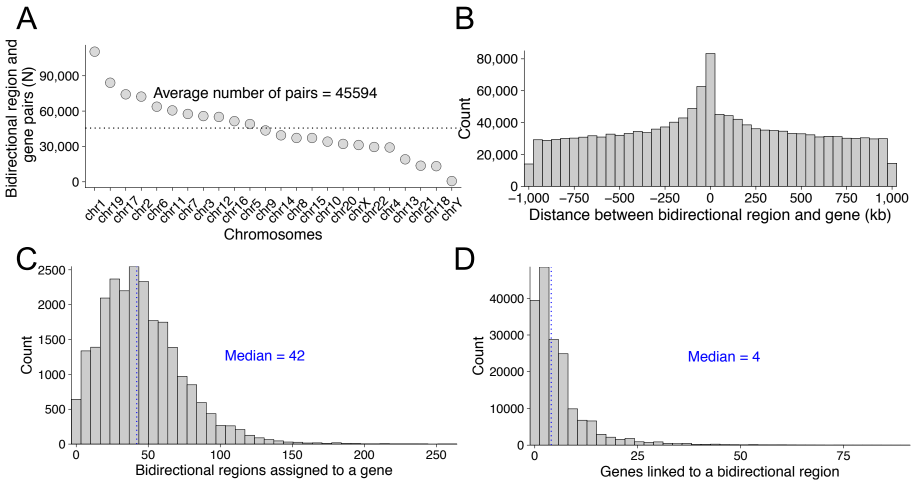


Figure 18: **Summary of gene and bidirectional pairs identified by the tissue specific interactions.** (A) Number of gene and bidirectional pairs per chromosome in human samples. (B) Distribution of distance between pairs (distance from bidirectional center to TSS of a gene). (C) Distributions of the number of bidirectional regions assigned to a gene across all tissues. (D) Distributions of the number of genes linked to a bidirectional region across all tissues.

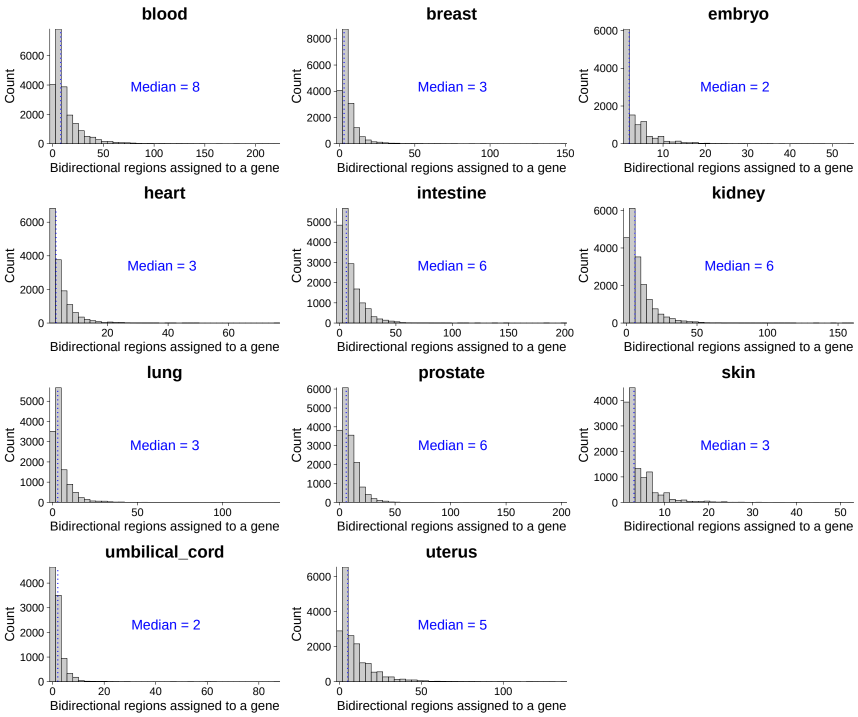


Figure 19: **Number of bidirectional regions assigned to a gene in each tissue.** Distribution of the number of bidirectional regions assigned to each gene in all 11 tissues assessed. The median is highlighted in each plot.

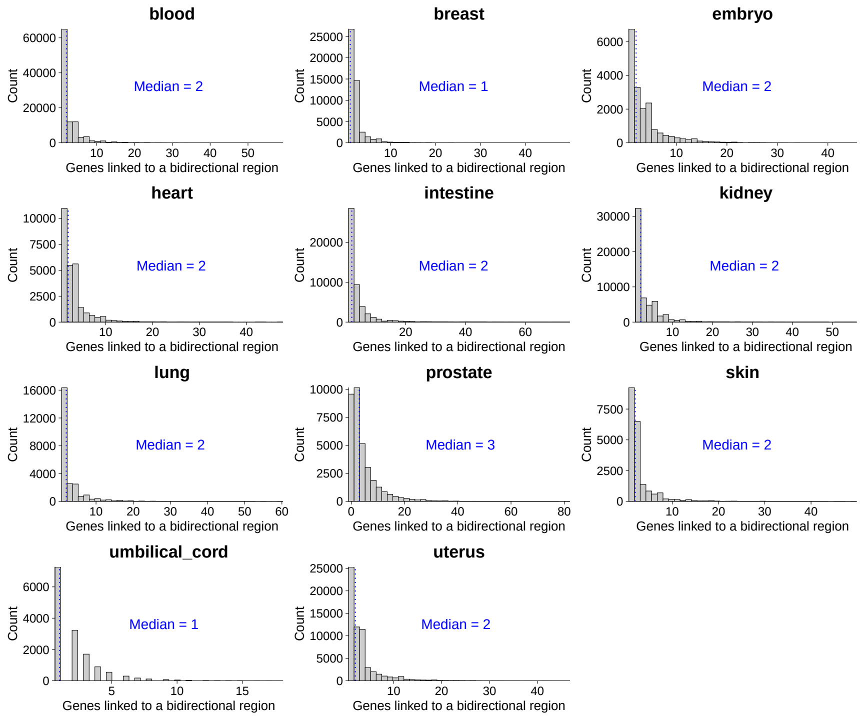


Figure 20: **Number of genes linked to a bidirectional region in each tissue.** Distribution of the number of gene transcripts linked to a bidirectional region in all 11 tissues assessed. The median is highlighted in each plot.

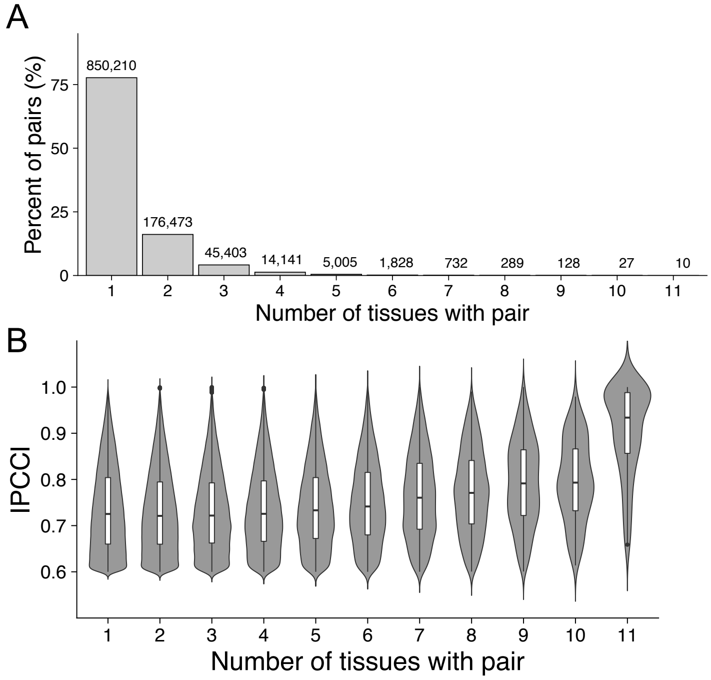


Figure 21: **Summary of correlated gene and bidirectional pairs identified in each tissue.** (A) The number and percent of *gene* $\rightarrow$ *bidirectional* pairs found in the tissues analyzed. (B) The |PCC| for gene and bidirectional pairs binned by the number of tissues they are found in.

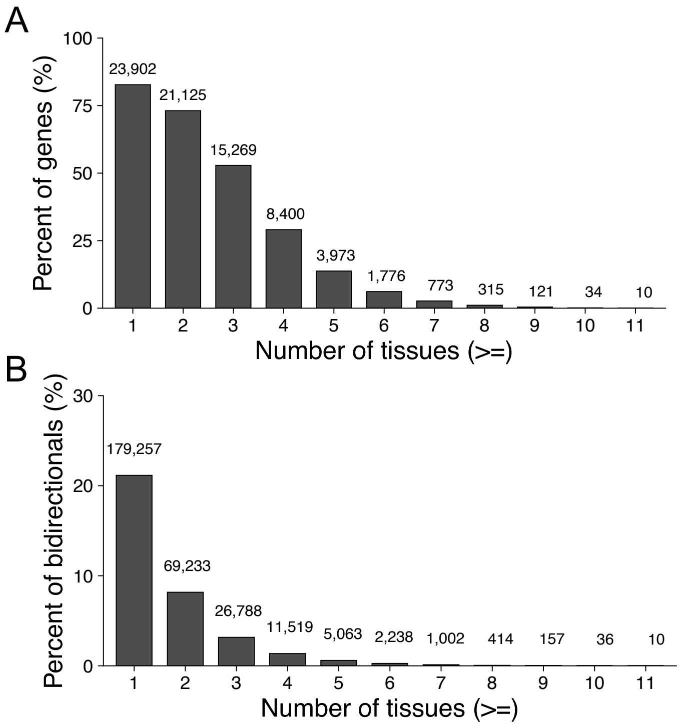


Figure 22: **Summary of gene and bidirectional regions used in the correlation analysis across all tissues.** (A) Number of genes assigned to a bidirectional region across all tissues. (B) Number of bidirectional regions linked to a gene transcripts across tissues.

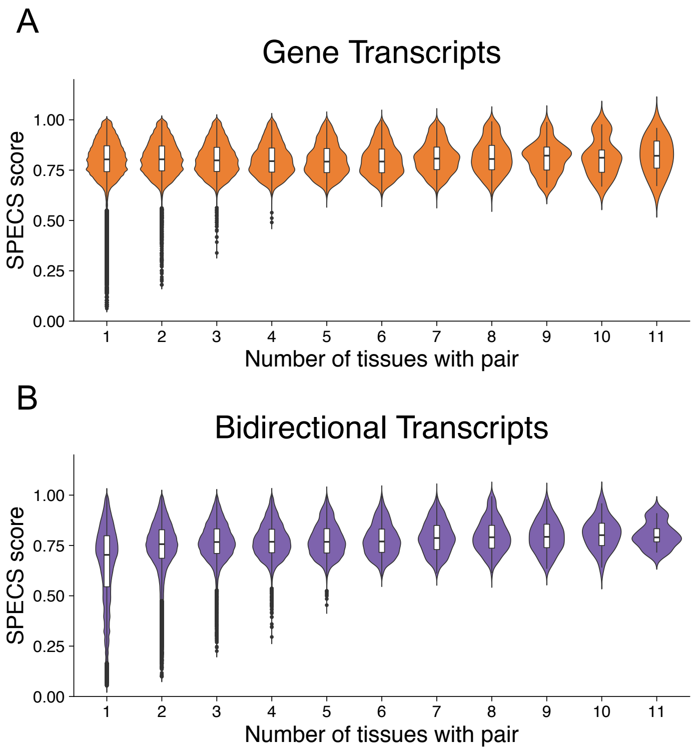


Figure 23: **SPECS scores for gene and bidirectional pairs identified in each tissue.** Maximum SPECS scores for (A) gene and (B) bidirectional region for significant pairs compared to the number of tissues they are identified in.

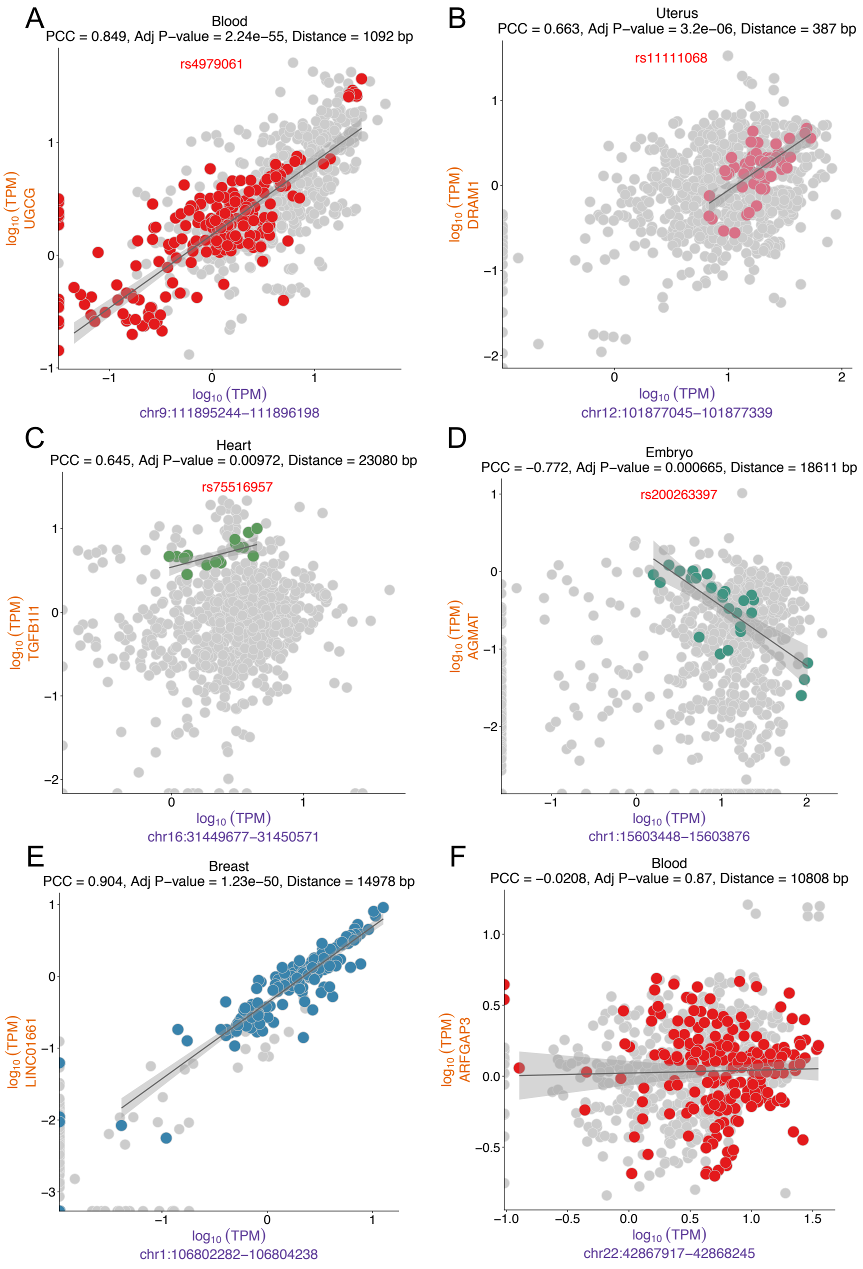


Figure 24: **Examples of gene and bidirectional pairs identified by the correlation method.** Continued on next page...

Figure 24: **Continued ...** Each point is a sample with colored dots highlighting the specified tissue (normalized counts for x-axis: bidirectional region and y-axis: gene). (A) UGCG in blood samples. The bidirectional region overlaps rs4979061. (B) DRAM1 and a bidirectional region in the promoter region found in uterus tissues. The bidirectional region overlaps rs11111068. (C) TGFB1I1 and bidirectional region found in heart tissues. Bidirectional region overlaps rs75516957. (D) AGMAT and bidirectional pair found in embryo samples, with bidirectional region overlapping rs200263397. (E) Significant LINC01661 and bidirectional pair found in breast samples. (F) Non-significant pair for ARFGAP3, shown for contrast.

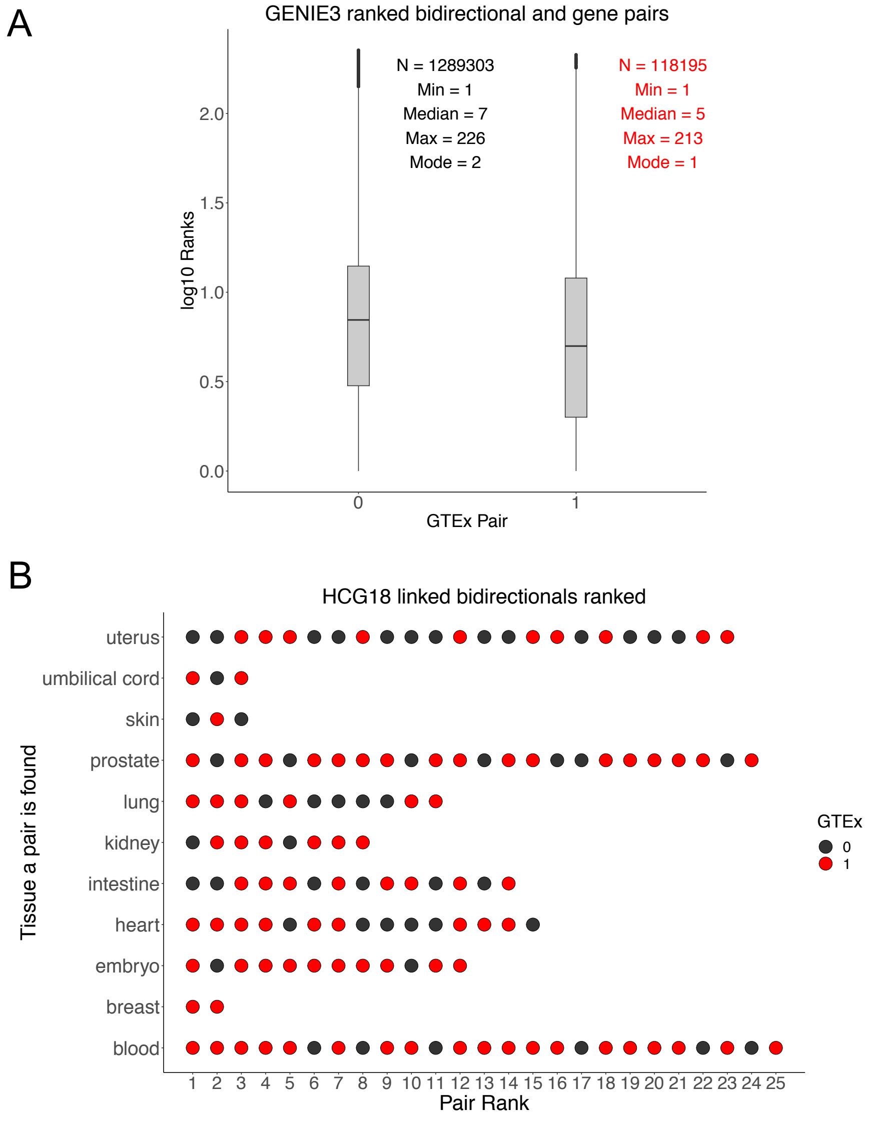


Figure 25: **GENIE3 ranks for correlated gene and bidirectional pairs.** (A) Box plot of pair ranks between those found in GTEx (1) compared to those not found in GTEx (0). Summary statistics for each category are also listed. (B) HCG18 and bidirectional regions ranked in each tissue a pair is found. The pairs found in GTEx are colored red. Blood pairs are shown in Figure 4E-F.

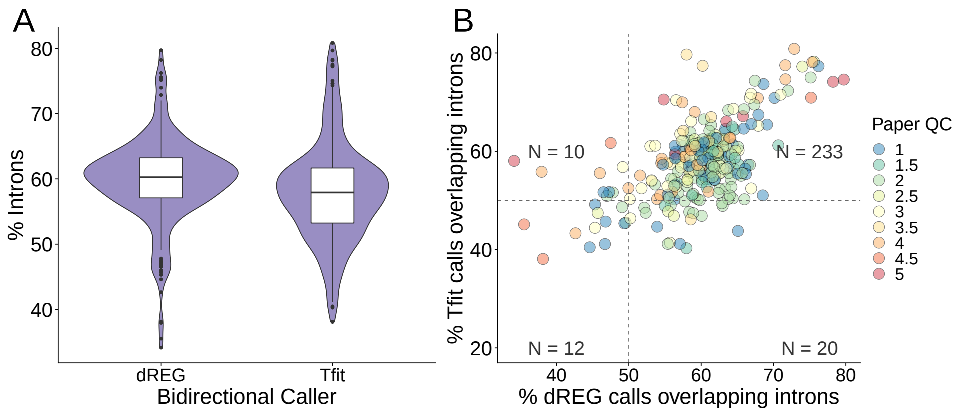


Figure 26: **Percent of paper bidirectional regions that overlap introns for dREG and Tfit calls.** (A) Distribution of percent intronic bidirectional regions called by dREG and Tfit after *muMerge* in each paper. (B) Scatterplot of dREG and Tfit intronic bidirectional regions in each paper across mouse and human samples. Each point is colored by the average paper QC. The vertical and horizontal lines are arbitrarily at 50%.

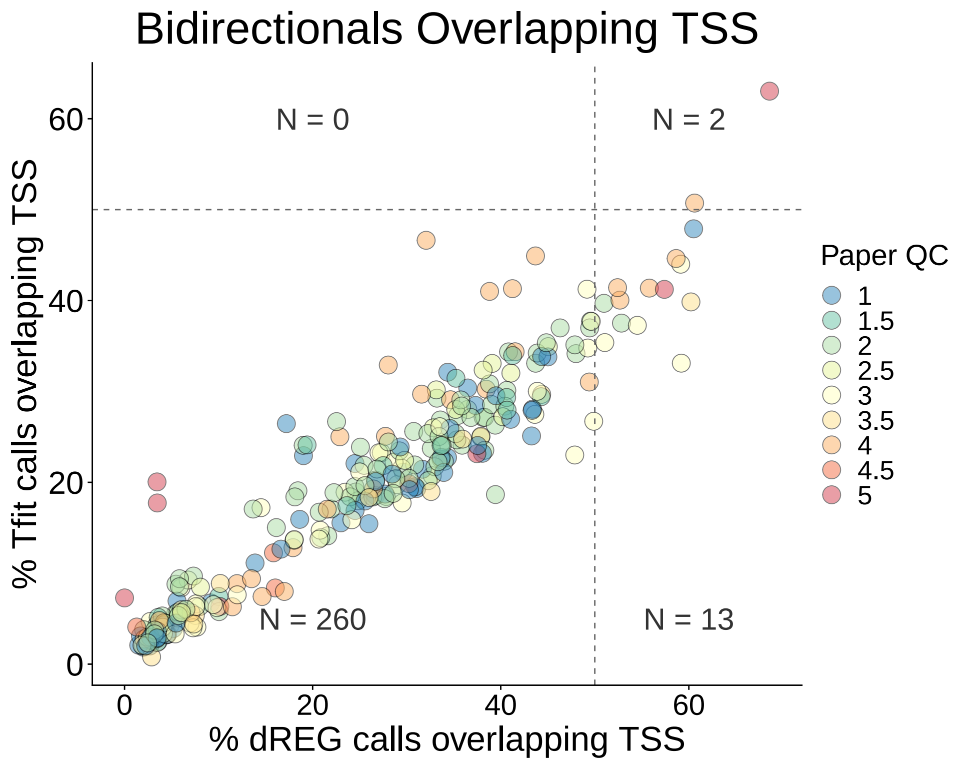


Figure 27: **Percent of called bidirectional regions that overlap annotated TSS, by paper.** Each point is colored by the average paper QC. The vertical and horizontal lines are arbitrarily at 50%.

### Supplementary Tables

| **Metadata Collected** | **Description** |
|---|---|
| Paper identifier | The paper where the samples and results were published, presented in the format *AuthorYearTitle_Word* (e.g. Allen2014global) |
| Sample identifier | The project identifier (SRP) and the sample identifier (SRR) |
| Replicate | Specify whether the sample was a technical or biological replicate |
| Organism | The scientific name of the organism (formatted as H. Sapiens or D. Melonogaster) |
| Genetic background | The cell type or tissue used |
| Modification | Genetic modifications such as RNAi, shRNA, if applicable |
| Treatment | The treatment, if applicable |
| Treatment times | Time the sample was exposed to a treatment |
| Nascent protocol | GRO-seq, PRO-seq, NET-seq, GRO-cap, PRO-cap, etc |
| Library preparation | Ligation, circularization, random primed and template switch reverse transcription |
| Spike in | If applicable, specify which spike-in control was used (ERCC, Drosophila, Arabopsis or Luciferase/Gal4/NeoR/GFP) |
| Tissue | The tissue type for each sample collected (e.g. adipose, blood, bone, brain, breast, embryo, eye, gonad, heart, inflorescence and meristem, intestine, kidney, leaf, liver, lung, micronucleus, multiple, muscle, nervous, ovary, pituitary, prostate, seed, seedling, shoot, skin, spleen, testis, umbilical cord, uterus and yolk sac) |

Table 1: Summary of the metadata manually collected for each dataset. Metadata collected from the original publication or from the corresponding GEO and/or SRA entry.

| | All Transcripts | Long Isoform |
|---|---|---|
| Total | 42,224 | 28,889 |
| Bidirectional region over TSS | 34,647 (82.1%) | 23,145 (80.1%) |

Table 2: Number of RefSeq gene transcripts with TSS bidirectional region in human samples.

| | 0 counts | <100 counts | <1000 counts | Median counts |
|---|---|---|---|---|
| With TSS (N=23,145) | 0.12% (28) | 1.23% (284) | 4.37% (1,012) | 430,404 |
| Without TSS (N=5,744) | 01.93% (111) | 15.86% (911) | 35.41% (2,034) | 3,628 |

Table 3: Transcription levels of long isoform (Ref-Seq transcripts) in human samples, with and without TSS associated bidirectional region identified.

| Tissue | SNPs | Bidirectional Regions | Genes |
|---|---|---|---|
| blood | 126 | 111 | 259 |
| breast | 75 | 65 | 144 |
| embryo | 25 | 21 | 69 |
| heart | 43 | 36 | 101 |
| intestine | 63 | 54 | 243 |
| kidney | 74 | 62 | 189 |
| lung | 30 | 26 | 106 |
| prostrate | 51 | 44 | 193 |
| skin | 30 | 20 | 76 |
| uterus | 84 | 70 | 233 |
| umbilical cord | 18 | 14 | 32 |

Table 4: Number of linked SNPs, bidirectional regions and genes in each tissue for the European GWAS catalog of leukemia associated SNPs (EFO_0000565).

[167] Link, V.M., Duttke, S.H., Chun, H.B., Holtman, I.R., Westin, E., Hoeksema, M.A., Abe, Y., Skola, D., Romanoski, C.E., Tao, J., Fonseca, G.J., Troutman, T.D., Spann, N.J., Strid, T., Sakai, M., Yu, M., Hu, R., Fang, R., Metzler, D., Ren, B., Glass, C.K.: Analysis of genetically diverse macrophages reveals local and domain-wide mechanisms

that control transcription factor binding and function. Cell **173**(7), 1796–180917 (2018)

[168] Linna-Kuosmanen, S., Tomas Bosch, V., Moreau, P.R., Bouvy-Liivrand, M., Niskanen, H., Kansanen, E., Kivelä, A., Hartikainen, J., Hippeläinen, M., Kokki, H., Tavi, P., Levonen, A.-L., Kaikkonen, M.U.: Nrf2 is a key regulator of endothelial microrna expression under proatherogenic stimuli. Cardiovascular Research (2020)

[175] Liu, W., Duttke, S.H., Hetzel, J., Groth, M., Feng, S., Gallego-Bartolome, J., Zhong, Z., Kuo, H.Y., Wang, Z., Zhai, J., Chory,

J., Jacobsen, S.E.: RNA-directed DNA methylation involves co-transcriptional small-RNA-guided slicing of polymerase V transcripts in Arabidopsis. Nature Plants **4**(3), 181–188 (2018)

[176] Liu, M., Zhu, J., Dong, Z.: Immediate transcriptional responses of arabidopsis leaves to heat shock. Journal of Integrative Plant Biology **n/a**(n/a) (2020)

[183] Magnuson, B., Veloso, A., Kirkconnell, K.S., de Andrade Lima, L.C., Paulsen, M.T., Ljungman, E.A., Bedi, K., Prasad, J., Wilson, T.E.,

Ljungman, M.: Identifying transcription start sites and active enhancer elements using BruUV-seq. Scientific reports **5** (2015)

[199] Mukai, C., Choi, E., Sams, K.L., Klampen, E.Z., Anguish, L., Marks, B.A., Rice, E.J., Wang, Z., Choate, L.A., Chou, S.-P., Kato, Y., Miller,

A.D., Danko, C.G., Coonrod, S.A.: Chromatin run-on sequencing analysis finds that ecm remodeling plays an important role in canine hemangiosarcoma pathogenesis. BMC Veterinary Research **16**(1), 206 (2020)

[248] Stender, J.D., Nwachukwu, J.C., Kastrati, I., Kim, Y., Strid, T., Yakir, M., Srinivasan, S., Nowak, J., Izard, T., Rangarajan, E.S., Carlson, K.E., Katzenellenbogen, J.A., Yao, X.-Q., Grant, B.J., Leong, H.S., Lin, C.-Y., Frasor, J., Nettles, K.W., Glass, C.K.: Structural

and molecular mechanisms of cytokine-mediated endocrine resistance in human breast cancer cells. Mol Cell **65**(6), 1122–11355 (2017)

[255] Tan, J.L., Fogley, R.D., Flynn, R.A., Ablain, J., Yang, S., Saint-André, V., Fan, Z.P., Do, B.T., Laga, A.C., Fujinaga, K., Santoriello, C., Greer, C.B., Kim, Y.J., Clohessy, J.G., Bothmer, A., Pandell, N.,

Avagyan, S., Brogie, J.E., van Rooijen, E., Hagedorn, E.J., Shyh-Chang, N., White, R.M., Price, D.H., Pandolfi, P.P., Peterlin, B.M., Zhou, Y., Kim, T.H., Asara, J.M., Chang, H.Y., Young, R.A., Zon, L.I.: Stress from nucleotide depletion activates the transcriptional regulator HEXIM1 to suppress melanoma. Molecular Cell **62**(1), 34–46 (2016)

[278] Wang, L., Zhou, Y., Xu, L., Xiao, R., Lu, X., Chen, L., Chong, J., Li, H., He, C., Fu, X.-D., Wang, D.: Molecular basis for 5-

carboxycytosine recognition by RNA polymerase II elongation complex. Nature **523**(7562), 621–5 (2015)

[286] Wei, C., Qiu, J., Zhou, Y., Xue, Y., Hu, J., Ouyang, K., Banerjee, I., Zhang, C., Chen, B., Li, H., Chen, J., Song, L.-S., Fu, X.-D.: Repression of the central splicing regulator RBFox2 is functionally linked to pressure overload-induced heart failure. Cell Rep (2015)

[296] Xi, Y., Shi, J., Li, W., Tanaka, K., Allton, K.L., Richardson, D., Li, J., Franco, H.L., Nagari, A., Malladi, V.S., Coletta, L.D., Simper, M.S., Keyomarsi, K., Shen, J., Bedford, M.T., Shi, X., Barton, M.C., Lee Kraus, W., Li, W., Dent, S.Y.R.: Histone modification profiling in

breast cancer cell lines highlights commonalities and differences among subtypes. BMC Genomics **19**(1), 150 (2018)

[307] Zhang, J., Cavallaro, M., Hebenstreit, D.: Timing polymerase pausing with TV-PRO-seq. bioRxiv (2018)

[308] Zhang, Z., Chng, K.R., Lingadahalli, S., Chen, Z., Liu, M.H., Do, H.H., Cai, S., Rinaldi, N., Poh, H.M., Li, G., Sung, Y.Y., Heng, C.L., Core, L.J., Tan, S.K., Ruan, X., Lis, J.T., Kellis, M., Ruan, Y., Sung, W.-K., Cheung, E.: An AR-ERG transcriptional signature defined by long-range chromatin interactomes in prostate cancer cells. Genome Research **29**(2), 223–235 (2019)

[309] Zhang, X., Zhang, Y., Ba, Z., Kyritsis, N., Casellas, R., Alt, F.W.: Fundamental roles of chromatin loop extrusion in antibody class switching. Nature **575**(7782), 385–389 (2019)

[310] Zhang, M., Lai, Y., Krupalnik, V., Guo, P., Guo, X., Zhou, J., Xu, Y., Yu, Z., Liu, L., Jiang, A., Li, W., Abdul, M.M., Ma, G., Li, N., Fu, X., Lv, Y., Jiang, M., Tariq, M., Kanwal, S., Liu, H., Xu, X., Zhang, H., Huang, Y., Wang, L., Chen, S., Babarinde, I.A., Luo, Z., Wang, D., Zhou, T., Ward, C., He, M., Ibañez, D.P., Li, Y., Zhou, J., Yuan, J., Feng, Y., Arumugam, K., Di Vicino, U., Bao, X., Wu, G., Schambach, A., Wang, H., Sun, H., Gao, F., Qin, B., Hutchins, A.P., Doble, B.W., Hartmann, C., Cosma, M.P., Qin, Y., Xu, G.-L., Chen, R., Volpe, G., Chen, L., Hanna, J.H., Esteban, M.A.: $\beta$-catenin safeguards the ground state of mouse pluripotency by strengthening the robustness of the transcriptional apparatus. Science Advances **6**(29) (2020)

[311] Zhang, X., Yoon, H.S., Chapdelaine-Williams, A.M., Kyritsis, N., Alt, F.W.: Physiological role of the 3'igh cbes super-anchor in antibody class switching. Proceedings of the National Academy of Sciences **118**(3) (2021)

[312] Zhao, Y., Liu, Q., Acharya, P., Stengel, K.R., Sheng, Q., Zhou, X., Kwak, H., Fischer, M.A., Bradner, J.E., Strickland, S.A., Mohan,

S.R., Savona, M.R., Venters, B.J., Zhou, M.-M., Lis, J.T., Hiebert, S.W.: High-resolution mapping of RNA polymerases identifies mechanisms of sensitivity and resistance to BET inhibitors in t(8;21) AML. Cell Reports **16**(7), 2003–2016 (2016)
